# Network dynamics scale with levels of awareness

**DOI:** 10.1101/2021.04.12.439452

**Authors:** Peter Coppola, Lennart R.B. Spindler, Andrea I. Luppi, Ram Adapa, Lorina Naci, Judith Allanson, Paola Finoia, Guy B. Williams, John D. Pickard, Adrian M. Owen, David K. Menon, Emmanuel A. Stamatakis

## Abstract

Small world topologies are thought to provide a valuable insight into human brain organisation and consciousness. However, functional magnetic resonance imaging studies in consciousness have not yielded consistent results. Given the importance of dynamics for both consciousness and cognition, here we investigate how the diversity of brain dynamics pertaining to small world topology (quantified by sample entropy; dSW-E) scales with decreasing levels of awareness (i.e., sedation and disorders of consciousness). Paying particular attention to result reproducibility, we show that dSW-E is a consistent predictor of levels of awareness even when controlling for the underlying functional connectivity dynamics. We find that dSW-E of subcortical and cortical areas are predictive, with the former showing higher and more robust effect sizes across analyses. Consequently, we propose that the dynamic reorganisation of the functional information architecture, in particular of the subcortex, is a characteristic that emerges with awareness and has explanatory power beyond that of the complexity of dynamic functional connectivity.

## INTRODUCTION

Recent neuroscience endeavours have approached the intractable question of consciousness via notions of complexity (Carhart-Harris et al., 2014; Northoff & Huang, 2017; Tononi, Boly, Massimini, & Koch, 2016; Varley et al., 2020). A complex system can be defined as a large network of components that exhibit collective emergent properties (Mitchell, 2011). In fact, consciousness researchers have focused their attention not only on the activity of brain regions; but also the statistical relationship between them (i.e., “connectivity”) and the resulting emergent global properties (Di Perri et al., 2016; Edelman & Gally, 2013; Stamatakis, Adapa, Absalom, & Menon, 2010; Tononi et al., 2016).

A prominent paradigm to investigate the complexity of brain connectivity is given by network science(Rubinov & Sporns, 2010; Watts & Strogatz, 1998). Using the mathematical framework of Graph Theory Analysis (GTA), network science permits an investigation into the topological/architectural characteristics of a network by defining its components as nodes and their interactions as edges. Watts and Strogatz in 1998 (Watts & Strogatz, 1998) brought this approach to the forefront by showing that complex real-life networks of the most disparate kinds tend to show a “small-world” (SW) architecture. Computationally, the SW network structure can be created by taking a regular lattice network (where neighbouring nodes are connected) and randomly rewiring some edges. This particular network configuration simultaneously retains many clusters of connected nodes, whilst the rewired edges enable information to travel easily across long distances in the network (i.e., an average “short path length”). The SW network is appealing to neuroscience as it putatively describes the fundamental local-global interaction of a limited number of brain regions and connections, and thus would allow complexity to emerge in a cost-effective manner (Bassett & Bullmore, 2017; Northoff & Huang, 2017; Sporns & Zwi, 2004; van den Heuvel, Stam, Boersma, & Hulshoff Pol, 2008). In fact, SW topology has been shown to favour synchronisation, richness of possible states, self-organisation, criticality, resistance to insult, efficient and cost-effective information transfer (Barahona, Barahona, & Pecora, 2002; Papo, Zanin, Martínez, & Buldú, 2016; Takagi, 2018, 2020; Tan & Cheong, 2017).

Given these characteristics, theorists have conjectured that SW organisation is relevant to consciousness (Alkire, Hudetz, & Tononi, 2008; Buzsáki, 2007; Carhart-Harris & Friston, 2019; Northoff & Huang, 2017; Sporns & Zwi, 2004). SW is a topology (i.e., interrelation of constituent parts) that indicates simultaneous localised-clustered (segregated) and efficient-global (integrated) information flow (Bassett & Bullmore, 2017; Deco, Tononi, Boly, & Kringelbach, 2015; Lord, Stevner, Deco, & Kringelbach, 2017; Northoff & Huang, 2017; Rubinov & Sporns, 2010), which theoretically would underpin the emergence of consciousness (Baars, 2005; Dehaene & Christen, 2011; Northoff & Huang, 2017; Sporns & Zwi, 2004; Tononi et al., 2016). In fact, SW has been theorised to underly the spatial temporal characteristics necessary for the emergence of awareness (Alkire et al., 2008; Buzsáki, 2007; Northoff & Huang, 2017).

The empirical side, conversely, has proposed several measures of SW architecture (Humphries & Gurney, 2008; Muldoon, Bridgeford, & Bassett, 2016; Telesford, Joyce, Hayasaka, Burdette, & Laurienti, 2011); but has not yielded the same level of consistency as its theoretical counterpart. Research in the functional network SW of anaesthesia has shown increases in SW during unconsciousness, in opposition to what would have been expected from theory (Monti et al., 2013; Northoff & Huang, 2017; Schroter et al., 2012). Others show decreases in SW during anaesthesia and disorders of consciousness (Barttfeld et al., 2015; Luppi et al., 2019). Still more papers report inconclusive SW results in consciousness-relevant conditions (Achard et al., 2012; Crone et al., 2014; Godwin, Barry, & Marois, 2015). There are also contradicting results arising from structural connectivity measurements of SW configurations (Tan et al., 2019; Weng et al., 2017).

Analogously to the proposed importance of small world topology to dynamic information flow (Barahona et al., 2002; Bassett & Bullmore, 2017; Takagi, 2018; Tan & Cheong, 2017; Watts & Strogatz, 1998), different theories of consciousness converge in proposing that the dynamic richness of possible brain states is a fundamental hallmark of consciousness (Carhart-Harris et al., 2014; Dehaene & Christen, 2011; Northoff & Huang, 2017; Tononi et al., 2016). This approach has in fact proven empirically successful (Barttfeld et al., 2015; Cavanna, Vilas, Palmucci, & Tagliazucchi, 2018; Demertzi et al., 2019; Golkowski et al., 2019; Huang, Zhang, Wu, Mashour, & Hudetz, 2020; Luppi et al., 2019).

Although SW is universally recognised as important for network dynamics (Barahona et al., 2002; Takagi, 2018; Tan & Cheong, 2017; Watts & Strogatz, 1998), SW studies of consciousness have primarily focused on static networks, created by averaging across time points (Achard et al., 2012; Crone et al., 2014; Monti et al., 2013; Schroter et al., 2012).

To tackle the inconsistencies between different empirical studies and theory, and to probe the relevance of network science to consciousness, we investigated the **dynamics** of small-worldness (SW). Specifically, we use an information-theory measure adapted to biological dynamical systems, namely **sample entropy** (Delgado-Bonal & Marshak, 2019; Richman & Moorman, 2000), to investigate how complex (“unpredictable” or “uncompressible”) SW architecture is over time. Given previous inconsistencies in this area, we devote particular attention to convergence of SW results by deploying different brain parcellations (i.e. region definitions that form network nodes). Parcellations, which varied between the aforementioned SW studies (Luppi et al., 2019; Monti et al., 2013; Schroter et al., 2012), have been known to affect graph theory results (Hallquist & Hillary, 2018; Luppi & Stamatakis, 2020; Papo et al., 2016; Yao, Hu, Xie, Moore, & Zheng, 2015). Therefore, the employment of different parcellations to assess whether results are parcellation-dependent is advised (Hallquist & Hillary, 2018). We used whole-brain parcellations with different granularities (i.e., Low and high granularity, 126 and 553 brain regions respectively, described in Supplementary material 1) and the AAL (Automatic Anatomical Labelling atlas), which has been extensively used in previous literature (Luppi et al., 2019; Schroter et al., 2012; Tan et al., 2019; Weng et al., 2017). To further assess convergence of results we chose to employ two different SW measures: Sigma (σ), as it is the most widely reported measure in the literature (Luppi et al., 2019; Monti et al., 2013; Schroter et al., 2012), and the more recently developed PHI (ϕ), which displays higher reliability in simulated networks and is designed for biologically-relevant weighted connectivity (Muldoon et al., 2016; Luppi 2021). Please note this is not “PHI” as defined in the context of integrated information theory (Tononi et al., 2016).

The empirical data consists of three independent functional magnetic resonance imaging (fMRI) datasets that are relevant to consciousness. Two are propofol anaesthesia datasets; the first collected in Cambridge (referred to as “CAM” dataset onwards), UK (18 participants) comprising a control awake and a moderate sedation condition (Adapa, Davis, Stamatakis, Absalom, & Menon, 2014) and the second in London, Ontario (henceforth referred to as LON) :16 participants in control awake and deep sedation conditions (Naci et al., 2018). The third dataset was acquired from patients with disorders of consciousness (hereafter indicated by “DOC”, Cambridge, UK). This comprised 23 patients of whom 11 are in a Minimally Conscious State (MCS), and the other 12 displaying the Unresponsive Wakefulness Syndrome (UWS).

These datasets are ideally suited to assess the importance of SW dynamics in consciousness as they permit an investigation which is independent of the type of consciousness alteration (i.e. pharmacologically or pathologically induced) and can be ordered according to decreasing levels of awareness (i.e., “content consciousness” (Laureys, Perrin, & Brédart, 2007)). We predict that the temporal complexity of SW, if relevant to consciousness, will consistently diminish with decreasing levels of awareness, in accordance to theoretical models (Carhart-Harris et al., 2014; Laureys et al., 2007; Tononi et al., 2016). If the sample entropy of small-world dynamics is consistently predictive of levels of awareness at the whole-brain level, we will investigate whether these effects are differentially driven by different subsystems (Cortex, Subcortex, Cerebellum). We will also test whether any subsystem effects are exclusive to SW or can be extended to other graph-theory properties that are relevant to consciousness in terms of segregation (functional division/specialisation) and integration (functional combination/information merging) (Achard et al., 2012; Luppi et al., 2019; Rubinov & Sporns, 2010; Sporns & Zwi, 2004).

## RESULTS

In order to confirm that the dynamics of network SW architecture can predict altered levels of awareness, we divided whole brain resting-state fMRI data spatially into different parcellations (Fig 1A) and then split the resulting timeseries using a sliding window approach (Barttfeld et al., 2015; Luppi et al., 2019; Preti, Bolton, & Van De Ville, 2017) (Fig 1B). Within each of these windows we constructed a network by relating each brain region’s timeseries to all others, using Pearson’s correlation coefficients (Fig 1C). Two small world measures (*PHI* (Muldoon et al., 2016); and *Sigma* (Humphries & Gurney, 2008), Fig 1D) were calculated on each of these networks. In this manner, we obtained a time-series of SW values on which sample entropy was calculated (Richman & Moorman, 2000) (Fig 1E), thus obtaining one value for each subject that denotes the richness, or complexity, of their SW fluctuations. Inferential statistics were performed using ordinal logistic regressions, with sample entropy values as the predictor variable and ordered conditions as the predicted variable (Fig 1F).

**Figure 1.**
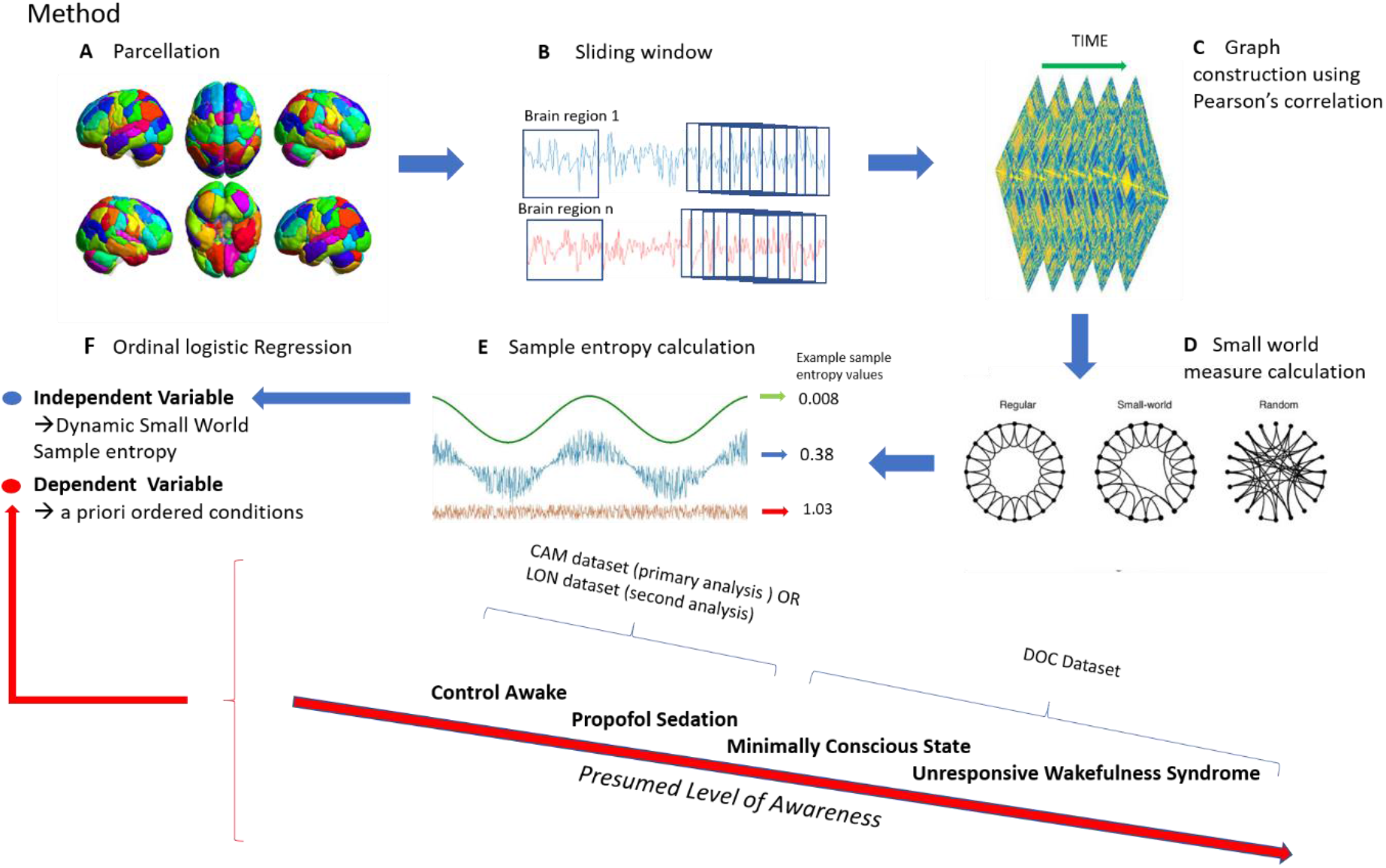
Method Description

To assess whether dynamic SW complexity scales with consciousness, we ordered the conditions *a-priori* according to presumed levels of awareness, analogously to previous studies looking at the functional network properties of consciousness (Demertzi et al., 2019; Di Perri et al., 2016). For the **main analysis** the CAM awake condition was ordered (as a factor in R) before the CAM moderate sedation, which in turn was placed as more aware than the DOC Minimally Conscious State (MCS) and DOC Unresponsive Wakefulness Syndrome (UWS) respectively. To assess the robustness of our results, we performed a **second analysis** by substituting the CAM propofol dataset with the LON propofol dataset which comprised an awake and a deep sedation condition (ordered respectively, Fig 1F).

### Illustration of method

A) We obtained timeseries for each brain region. B) The timeseries were divided with a sliding window approach (each window comprising of 24 timepoints and slided by 1 timepoint). C) We then correlated all region timeseries to obtain a weighted graph for each window. D) We calculated SW (Humphries & Gurney, 2008; Muldoon et al., 2016; Watts & Strogatz, 1998) for each graph so as to obtain a timeseries of SW values on which E) we calculated sample entropy. F) We inserted the sample entropy of dynamic small worldness into an ordinal logistic regression as a predictor; with the ordered conditions (according to presumed level of awareness) as a dependent variable.

#### SW dynamic complexity in the brain

Our hypothesis that dynamic SW sample entropy (dSW-E) predicts monotonically decreasing levels of awareness was confirmed using an **ordinal logistic regression** for both SW measures (Fig 2: **PHI** Standardized Regression Coefficients [Coef]= −1.42 p = 0.000006; C.I. [2.5%:97.5%] −2.05 : −0.81; **Sigma** Coef: −0.91 p=0.0002 C.I. −1.43 : −0.39). This result was corroborated across different parcellations with different granularities (presented in supplementary material 2). Furthermore, this result was replicated in the second analysis with a different sedation dataset (i.e., LON-DOC datasets= PHI; Coef= −0.82 p =0.001 C.I. −1.37 : −0.26 and Sigma; Coef = −0.81 p =0.001, C.I. −1.32 : −0.29; S2, Fig2). This suggests that the unpredictability of dynamic SW architecture reliably scales with increasing levels of awareness.

**FIGURE 2.**
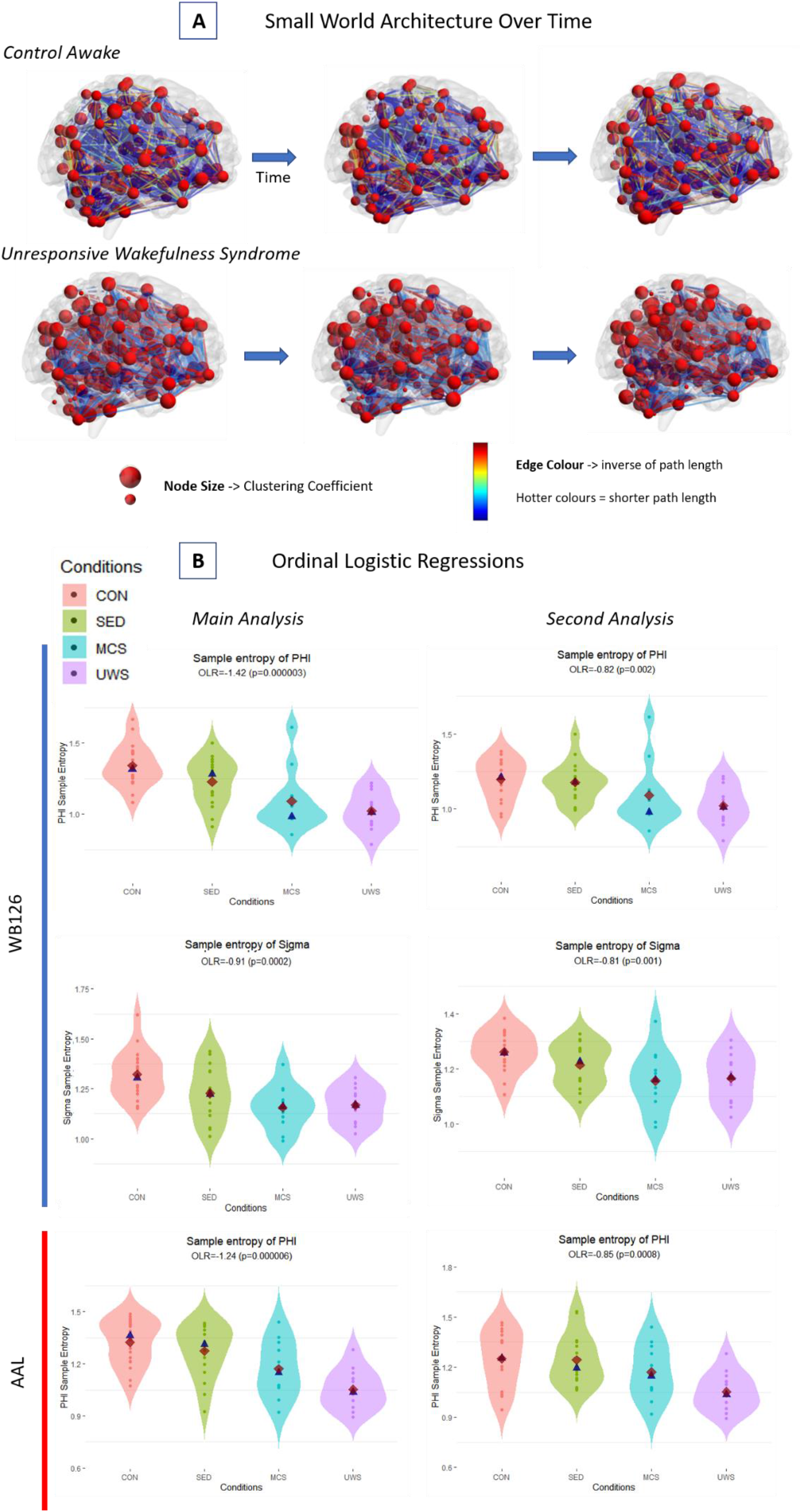
Dynamic SW properties and Ordinal Logistic Regression results.

This consistency is remarkable given that when we calculated the two SW measures (Phi and Sigma) on static graphs (i.e. one graph per participant constructed by averaging across all timepoints (Monti et al., 2013; Schroter et al., 2012)), they were not correlated (Rho=0.17, p=0.2) and did not yield consistent patterns between conditions and network definitions (S3). Conversely, the two measures of SW when calculated dynamically, proved more informative and showed the same intuitive patterns of decreasing complexity in lower levels of awareness (Fig 2, S2).

It is important to assess whether these graph theory entropy metrics truly reflect the temporal complexity of the functional architecture (i.e., topology), or can be explained more parsimoniously by lower order metrics such as the variation in functional connectivity (FC) (van den Heuvel et al., 2017). In fact, the entropy of average positive dynamic FC (chosen as graph theory properties are here calculated on positive correlations (Rubinov & Sporns, 2010; van den Heuvel et al., 2017) was a significant predictor for levels of awareness across parcellations and datasets (e.g., lower granularity whole brain parcellation for CAM-DOC analysis: Coef=−0.49, C.I.= −0.95:-0.02, p=0.01, see S4 for all results). Such results suggest that awareness entails an unpredictability of dynamic global synchronisation levels (measured by brain region timeseries Pearson’s correlations). This begs the question whether dynamic SW entropy (dSW-E, Fig 2) truly reflects the consciousness-predictive complexity of functional topology, or whether SW entropy results may be better explained by the unpredictability of global synchronisation.

To investigate this question, we ran a control ordinal logistic regression analysis that involved the same exact procedure used above (fig 1f) with the addition of the sample entropy of dynamic FC (dFC-E) as a covariate predictor. Both SW entropy predictors remained significant in the main analysis (S5). However, in the analysis controlling for dFC-E in the LON-DOC dataset, the dSW-E of Sigma for the AAL parcellation lost significance (S5), whilst dSW-E results remained significant in all other parcellations despite controlling for dFC-E.

This control analysis included both dFC-E and dSW-E as co-variate predictors in the same ordinal logistic regression and found the latter remained significant. This suggests that the temporal complexity of the functional SW architecture predicts increasing levels of awareness above and beyond what can be explained by the complexity (“compressibility”) of dynamic functional connectivity. This may be taken as a strong indication that the dynamic information produced (measured via sample entropy(Richman & Moorman, 2000) by functional topological dynamic organisation will decrease with diminishing levels of awareness.

In the top panel **(A)** are shown example dynamic SW properties (3 timepoints) overlaid on a brain template (Brainmesh_Ch2withCerebellum; BrainNet viewer(Xia, Wang, & He, 2013)]). The size of the nodes represents the clustering coefficient of that node, whilst the connections represent the inverse of the path length between the two nodes. The hotter the colour, the shorter the path length (i.e., how easily information can be transmitted between the nodes, not direct connectivity). Shown, for illustrative purposes, (top-left) is an example of a control awake participant (CAM dataset) and an example of an Unresponsive Wakefulness Syndrome patient (DOC dataset; bottom-left). Noticeable is the change in node size (clustering coefficient) and edge colour (path length) over time in the control participant, which is not so prominent in the individual affected by UWS. In the bottom panel **(B)** violin plots are showing the scaling of dynamic SW sample entropy with levels of awareness for both the main and second analysis. Conditions are ordered (left to right) according to a-priori presumed level of awareness (i.e. Awake > Propofol Sedation > MCS > UWS). The first two rows represent the dynamic PHI and Sigma entropy for a whole brain parcellation with 126 regions (S1) respectively. The third row represents dynamic PHI entropy values for the AAL. Blue triangle represents the median, whilst the red diamond represents the mean. OLR= Ordinal logistic regression coefficients; UWS= unresponsive wakefulness syndrome; MCS= minimally conscious state; SED = propofol sedation; Con= control awake condition. All ordinal logistic regression values are standardized for comparison.

#### Dynamic Small-world Entropy in the Cortex, Subcortex and Cerebellum

Given that dynamic entropy of both SW measures (φ & σ) consistently predicts levels of awareness at the whole-brain level, we sought to explore whether the dSW-E of major cito-architectonically distinct subdivisions of the brain (i.e., Cortex, subcortex and cerebellum) are relevant to consciousness and differentially explain the above whole-brain effects found in both high and low granularity parcellations.

This analysis is relevant to debates in the literature in which, some postulate the importance of the cortex in consciousness (Ledoux & Brown, 2017; Tononi et al., 2016); whilst others posit an essential role for the subcortex (Carhart-Harris & Friston, 2019; Panksepp, 2011; Solms, 2013). Although Sigma was used in the whole-brain analyses to assess convergence of different SW measures, and because it is widely used in published literature; in this section we exclusively calculated the sample entropy of dynamic PHI. We continued analyses with this particular measure because it is the most stable and computationally viable SW metric, specifically designed for weighed networks (Muldoon et al., 2016).

The complexity of dynamic SW topology predicted levels of awareness in the cortical (Coef= −1.30 p=0.000006 −1.89:-0.71) and subcortical (Coef=−1.94 p=0.000003 C.I.= −2.7:-1.10) network definitions. The cerebellar parcellation displays a significant trend (fig 3 Coef= −0.52, p=0.02, C.I.= −1.06:0.01).

**Figure 3.**
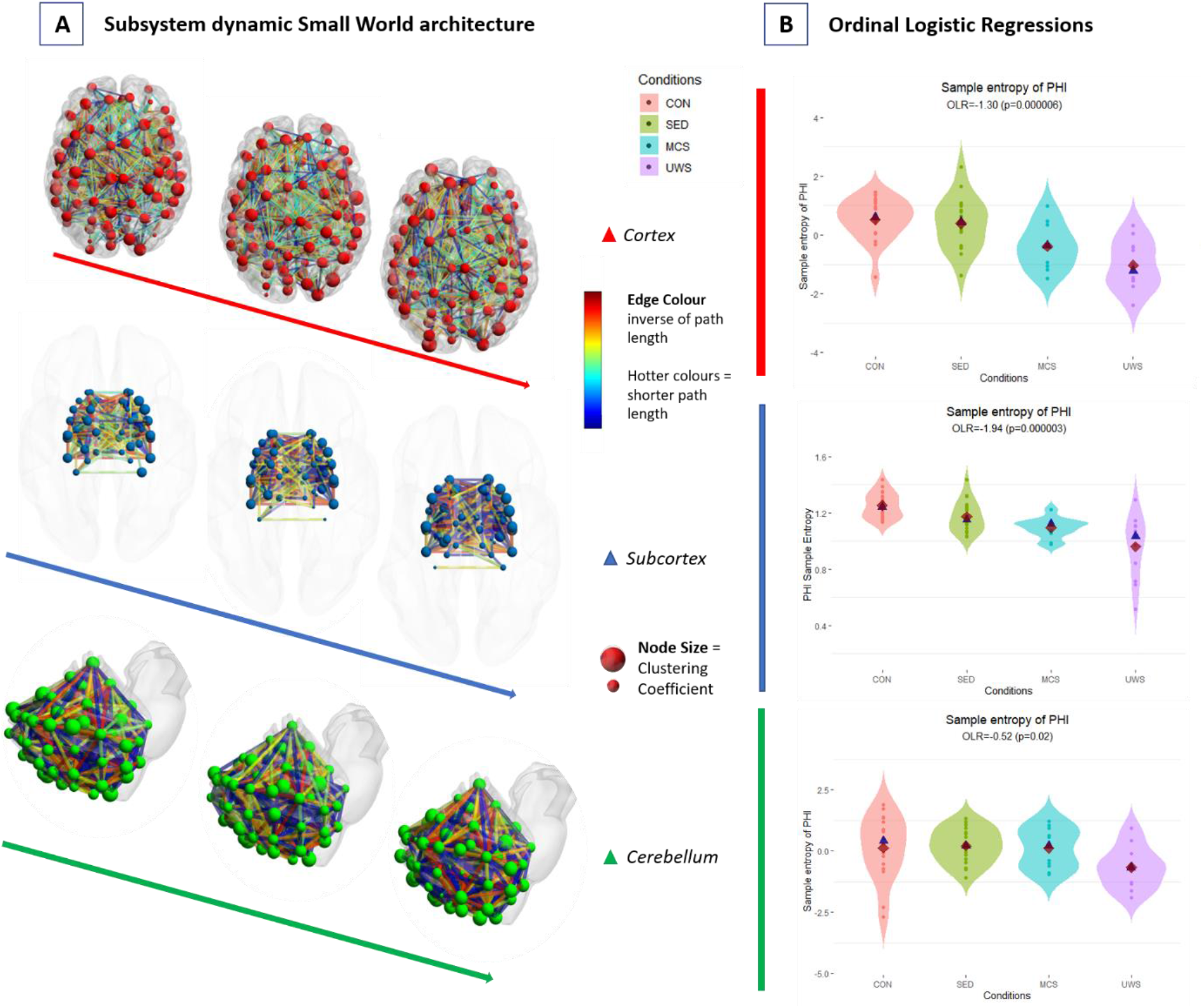
Subsystem dynamic SW properties and Ordinal Logistic Regression results.

Remarkably, the effect sizes for the subcortex were greater than those of the cortex. The LON-DOC datasets showed convergent results with the exception of the high granularity cortical parcellation (400 nodes; p=0.08, S6). Furthermore, when we added dFC-E as a covariate to dSW-E, both the cortex and subcortex remained significant, whilst the cerebellum dSW-E was borderline significant (p=0.055; S7). When controlling for dFC-E, the Subcortex dSW-E again had higher regression coefficients (coef=−1.83) than the cortex (coef=−1.25) in both datasets, although the cortex violated regression assumptions in the CAM-DOC analysis (S7).

To confirm whether the subcortex dSW-E was more predictive of levels of awareness than the cortex, we inserted dSW-E of the three subsystems within the same model and calculated the odds ratio for each. We found that an increase of 1 of the subcortical dSW-E increased the chance of being in the highest level of awareness by 6.34 (C.I.= 2.60:17.71), whilst a unit increase of cortical dSW-E increased the likelihood of being in the highest awareness category by 2.81 (C.I.= 1.50:5.52). Instead, the Odds ratio of the cerebellar dSW-E was 1.26 (C.I.= 0.70:2.32). A similar pattern was seen in the LON-DOC dataset (S8). These results suggest that the complexity of subcortical dynamic SW topology may be more consistently sensitive to decreasing levels of awareness than the cortex.

We then investigated whether these measures were correlated to behavioural metrics (bedside diagnostic assessments of consciousness-impairment in DOC patients via the Coma recovery Scale-revised [CRS-r (Giacino, Kalmar, & Whyte, 2004)]; and pharmacological plasma propofol concentration metrics (in CAM dataset). This would elucidate the potential relevance of dSW-E to clinically relevant observable behaviour (in the case of CRS-r) and the amount of propofol found in the blood. We found that subcortex dSW-E was inversely correlated to Propofol plasma concentration in the CAM dataset (rho= −0.38, p=0.022). However, this correlation did not survive Bonferroni correction for the different parcellations and measures used. There was also a correlation between SW entropy of PHI of the cortex and CRS-r (rho = 0.56, p=0.004), but this did not replicate in the higher granularity parcellation. Thus, the unpredictability of dynamic SW topology may be loosely related to observable behaviour and propofol concentration metrics.

In the left panel **(A)** are shown example dynamic SW properties (3 timepoints) for the cortex, subcortex and cerebellum ( templates: BrainMesh_Ch2withCerebellum; BrainMesh_ICBM152_smoothed; BrainMesh Cerebellum respectively, created in BrainNet viewer (Xia et al., 2013)). Node size represents the clustering coefficient of that node, whilst the connections represent the inverse of the path length between the two nodes. The hotter the colour, the shorter the path length. Shown are three timepoints for the cortex (red nodes; no=100), Subcortex (blue nodes; no=54) and the cerebellum (green nodes; no=99), for an awake participant (CAM dataset). In the right panel **(B)** violin plots are showing the scaling of dynamic PHI sample entropy with levels of awareness. Blue triangle represents the median, whilst the red diamond represents the mean. Conditions are ordered (left to right) according to a-priori presumed levels of awareness (i.e. Awake > Propofol Sedation > MCS > UWS). The first row shows the cortical, the second the subcortical and the last the cerebellar results. OLR= Ordinal logistic regression coefficients; UWS= unresponsive wakefulness syndrome; MCS= minimally conscious state; SED= propofol sedation; Con= control awake condition. All Ordinal Logistic Regression values are standardized for comparison.

#### Sample entropy of integration and segregation dynamic graph theory metrics for the cortex, subcortex and cerebellum

Given that subcortical dSW-E has more predictive power than the cortex, we sought to investigate whether this striking effect (Fig. 3) is specific to SW or can be generalised to other topological organisation (graph theory) measures that are relevant to consciousness theory. To this end, we chose two measures that are conceptually (in terms of integration and segregation (Lord et al., 2017; Rubinov & Sporns, 2010; Sporns & Zwi, 2004)) and statistically related to SW (Jarman, Steur, Trengove, Tyukin, & Van Leeuwen, 2017). The first is modularity (Q), which measures the extent to which a network can be divided, therefore being a proxy measure for segregation in terms of functional differentiation at a network level. The second measure is participation coefficient (PC), that measures to what extend different modules (i.e., functional subdivisions calculated via modularity) of the network are interconnected, therefore indicating the degree of integration in terms of the merging of information from different modules. We analysed these measures as we did SW in the previously described procedure (from Fig 1D onwards)

When we inserted the complexity of dynamic participation coefficient (dPC-E) of the different subsystems within the same model we saw that the odds ratio of the subcortex (2.14; C.I.=1.08:4.44, p=0.01) were higher than those that of the cortex (1.09; C.I.=0.63:1.90, p=0.01). Remarkably, the dPC-E of the cerebellum (2.76; C.I.=1.46:5.71, p=0.001) was the strongest predictor (S9). The complexity of modularity dynamics (dQ-E) had higher predictive power in the subcortex (2.15 C.I.=1.10:4.42, p=0.01) than the cortex (1.52; C.I.=0.86:2.75, p=0.07) and the cerebellum (1.61; C.I.=0.86:3.07, p=0.06). These results also reproduced in the second analysis (S9).

Using the behavioural scores collected in the original semantic propofol study for the CAM dataset (Adapa et al., 2014) we found that Subcortical dPC-E was inversely correlated to reaction time (Rho=−0.40, p=0.017), indicating the relevance of this measure to observable behaviour. These results suggest that decreasing subcortical complexity of network dynamics (beyond SW) is a characteristic of decreasing levels of awareness.

## DISCUSSION

In consonance with our hypothesis, a key finding of this study is that the temporal complexity of SW architecture increases with levels of awareness. We integrate graph theory (Achard et al., 2012; Crone et al., 2014; Monti et al., 2013; Schroter et al., 2012) and dynamics (Barttfeld et al., 2015; Demertzi et al., 2019; Huang et al., 2020; Luppi et al., 2019) in a novel way to show that the temporal complexity of information architecture reliably scales with levels of awareness. Importantly, we show that the complexity of subcortical dynamics is particularly predictive of levels of awareness. Such cortical and subcortical dynamics possibly underlie the varied streams of contents and states that characterise consciousness (Carhart-Harris & Friston, 2019; Dehaene & Christen, 2011; Panksepp, 2011; Solms, 2013; Tononi et al., 2016).

Previously, changes in static (Di Perri et al., 2016; Naci et al., 2018; Stamatakis et al., 2010) and dynamic functional connectivity (Barttfeld et al., 2015; Cavanna et al., 2018; Demertzi et al., 2019; Golkowski et al., 2019) in different states of consciousness have been widely reported. We advance this body of research by showing that the sample entropy of dynamic FC is predictive of levels of awareness; but importantly, we additionally show that SW architecture dynamics have consistent explanatory power above and beyond the variations in functional connectivity. This suggests that awareness has characteristic dynamic global information architectures (topologies) that cannot be reduced to simple FC. In other words, the dynamic re-configuration of the global functional architecture (“the interrelation of parts”), rather than the absolute synchronisation of brain regions, may be particularly important to consciousness. These findings, therefore, speak to theories that posit a global workspace (of information (Baars, 2005)), or the irreducibility of the whole to its parts (Tononi et al., 2016). In fact, we show that the dynamics of architectures that favour both integration and segregation between different information modules, consistently scale with increasing levels of awareness. It is therefore possible that such architectures may contribute to an integrated dynamic global workspace of information across time.

A key result of this study may supply some interpretations in regards to what may be particularly important for consciousness emergence. This is the difference between cortical and subcortical effects. Despite the cortex was a significant predictor on its own, we found that the complexity of dynamic subcortical topology is more consistent and powerful in predicting levels of awareness than the cortex. This suggests that the complexity of topological functional dynamics in the subcortex is particularly sensitive to different levels of awareness. In fact, the subcortical system is thought to provide fundamental (affective, interoceptive and sensory) inputs for cortical processing, and is hypothesised to have underpinned the first subjective experiences in evolutionary history and subsequent phylogenetic development of higher-order self-awareness (Carhart-Harris & Friston, 2019; Panksepp, 2011; Solms, 2013). Despite studies of the dynamics of consciousness tend to focus mainly on the cortex (Barttfeld et al., 2015; Demertzi et al., 2019; Huang et al., 2020), recent evidence (Lutkenhoff, Johnson, Casarotto, Massimini, & Monti, 2020) shows that the complexity of cortical response to perturbation in DOC inversely correlates with atrophy in arousal-related subcortical structures rather than in cortical structures.

Although some authors propose that subcortical structures function as an unconscious modulator of behaviour and cognitive conscious access (Ledoux & Brown, 2017), rather than underpinning basic awareness (Panksepp, 2011; Solms, 2013), these results suggest that formulations that focus on the cortex as central to consciousness (Baars, 2005; Tononi et al., 2016), may necessitate further verification in the future (e.g. see, Shewmon, Holmes, & Byrne, 1999). In fact, the present results suggest that investigating subcortical information may aid finer differentiation of different conscious states (Lutkenhoff, Johnson, et al., 2020; Lutkenhoff, Wright, et al., 2020; Panksepp, 2011) for clinical purposes.

In a similar vein, despite some researchers do not consider the Cerebellum important for consciousness (e.g., Tononi et al., 2016); we found that network dynamics of this subsystem (particularly dPC-E) displays some degree of predictive power for levels of awareness. This lends tentative support to notions that the cerebellum may have a discernible role in awareness (Clausi et al., 2017; Johnson, Belyk, Schwartze, Pinheiro, & Kotz, 2019).

Another contribution of this paper is the investigation of the relevance of SW as a metric for consciousness. Although several measures of SW have been proposed (Humphries & Gurney, 2008; Muldoon et al., 2016; Telesford et al., 2011); its calculation in static functional brain networks have been problematic (Rubinov & Sporns, 2010). Such issues, other than being evidenced by the inconsistency between previously published studies on consciousness (Achard et al., 2012; Barttfeld et al., 2015; Crone et al., 2014; Luppi et al., 2019; Monti et al., 2013; Schroter et al., 2012), were found within this study (S3). Here, conversely, we show that the richness of the dynamics of this topological measure robustly decreases with diminishing levels of awareness, independently of the cause of unconsciousness, dataset, brain region definition, and different measures of SW. The SW topology implies an information communication architecture that is simultaneously efficient and specialised. In fact, SW is thought to be related to information transmission and cognition in both health and disease (Achard, Salvador, Whitcher, Suckling, & Bullmore, 2006; Bassett & Bullmore, 2017; Schilling, 2005; Sporns & Zwi, 2004; Takagi, 2018; Tan & Cheong, 2017; van den Heuvel et al., 2008; Wu et al., 2012; Yu et al., 2011; Zhu et al., 2020). Therefore, it is possible that the reconfiguration of SW topology over time may indicate variations in information processing (and therefore cognitive states (Bassett & Bullmore, 2017)), which would intuitively increase proportionally to the level of awareness (Carhart-Harris et al., 2014; Laureys et al., 2007; Tononi et al., 2016).

However, we have also shown that consciousness-relevant topological dynamics are not limited to SW. The sample entropy of dynamic Modularity (Q) and PC, may index changes in the formation and inter-communication of dynamic functional subsystems (e.g., in visual attention), and as such may provide a good metric of variations in the stream of conscious contents or cognitive states that is typical for awareness (Di Perri et al., 2016; Dixon et al., 2017; Edelman & Gally, 2013; Godwin et al., 2015; Huang et al., 2020; Margulies et al., 2016). Either way, analogously to SW, PC and Q have been shown to be related to cognition and information processing (Arnemann et al., 2015; Bertolero, Yeo, Bassett, & D’Esposito, 2018; Cohen & D’Esposito, 2016; Finc et al., 2017; Godwin et al., 2015; Han, Chapman, & Krawczyk, 2020; Hilger, Ekman, Fiebach, & Basten, 2017). Thus, interpretations are complementary to those made above for SW, in that the dynamic entropy of these GTA properties may indicate variations in information processing state (and therefore contents of consciousness).

Given the potential existence of many different types (or dimensions) of consciousness, that the dynamic complexity of several graph theory properties may display predictive power, and that these measures display high within condition standard deviations (Fig 2 & 3); we tentatively suggest that these results may primarily relate to the epi-phenomenology of consciousness. In other words, the dynamic complexity of functional topology necessarily arises with consciousness, but it may not be a sufficient condition for the emergence of awareness.

As for the strengths and weakness of this study; the temporal resolution of the data-collection technique and the sliding window approach constitute a limitation of this study, as it only can measure coarse timescales of brain activity. Furthermore, DOC data is inherently noisy and is characterised by high degrees of variability and misdiagnosis. We selected this subset of participants out of a bigger dataset to ensure the data had acceptable quality. The ordering of conditions into decreasing levels of awareness may be controversial, in that it reduces subjective qualitative states to a two-dimensional quantity, despite being clinically (Giacino et al., 2004; Laureys et al., 2007), theoretically (Carhart-Harris et al., 2014; Tononi et al., 2016), and intuitively justified. Conversely, in light of inconsistencies between previously published studies, the use of different SW measures and related graph theory measures constitute a strength of this study. In fact, the explicit controlling for the dynamic FC (which underlies the graph theory measures) is a first in graph-theory consciousness research and consolidates the robustness and interpretation of results (van den Heuvel et al., 2017). The additional use of an independent dataset to validate results, and the use of different parcellations (with different brain region definitions but similar numbers, and with similar definitions but different granularities, S1) serve to augment assurance in these results (Hallquist & Hillary, 2018).

We conclude, with a reasonable amount of confidence, that the complexity of dynamic topology (in other words: the re-organisation of functional information architecture) does increase with the emerging of awareness. We tentatively suggest that dynamics of information processing architecture indirectly reflects changes in cognitive content/mental state which is an intuitive characteristic of the vernacular “stream of consciousness”. The predictive power of the subcortex’s dynamic topology is higher and more consistent compared to that of the cortex or the cerebellum, suggesting that the dynamic re-organisation of this system may be particularly important in typical awareness.

## Methods

### Cambridge anaesthesia dataset (CAM)

#### Participants – CAM dataset

Ethical approval was obtained from the Cambridgeshire 2 Regional Ethics committee (Adapa et al., 2014). 25 participants were recruited, however due to incomplete data in the cortex and procedure failure a subset of 18 were taken for further analyses. All participants were healthy and were native English speakers (50% males). Mean age was 33.3 (19-52). Two senior anaesthetists were present during scanning. Electrocardiography and pulse oximetry were continuously performed whilst measures of blood pressure, heart rate and oxygen saturation were recorded regularly.

#### Anaesthetic Protocol – Cam Dataset

Propofol sedation was administered intravenously via “target controlled infusion” with a Plasma Concentration mode. An Alaris PK infusion pump (Carefusion, Basingstoke, UK) was used which was controlled via the Marsh pharmacokinetic model. The anaesthesiologist can thus decide on a desired plasma 2 “target” and the system will regulate the infusion rates using patient characteristics as covariates. Three target plasma levels were used – no drug (awake control), 0.6 μg/ml (low sedation), 1.2 μg/ml (moderate sedation). In this study only the moderate sedation is used. Data for this latter condition was taken 20 minutes after cessation of sedation. Blood samples were taken at the end of each titration period, before plasma target was altered. The level of sedation was probed verbally immediately before and after each of the scanning runs.

10 minutes of plasma and effect-site propofol concentration equilibration was allowed before cognitive tests were commenced (auditory and semantic decision tasks). Mean (Standard deviation) plasma propofol concentrations was 304.8 (141.1) mg/ml during light sedation, 723.3 (320.5) mg/ml during moderate sedation and 275.8 (75.42) mg/ml during recovery. Mean (SD) total propofol given was 210.15 (33.16) mg.

#### Magnetic Resonance Imaging Protocol – Cam dataset

A Trio Tim 3 tesla MRI machine (Erlangen, Germany), with 12-channel head coil was used to obtain 32 descending interleaved oblique axial slices with an interslice gap of 0.75 mmm and an in plane resolution of 3 mm. The field of view was 192×192, Repetition time and acquisition time was 2 seconds whilst the echo time was 30 ms and flip angle 78. T1-weighted structural images with 1mm resolution were obtained using an MPRAGE sequeunce with TR= 2250 ms , TI– 900ms , TE= 2.99 ms flip angle= 9 degrees.

### London Ontario Propofol (LON) dataset

#### Participants-LON dataset

The second anaesthesia dataset used was obtained at the Robarts Research Institute in London, Ontario (Canada) and was approved by the Western University Ethics board. 19 healthy (13 males; 18-40 years), right-handed, English speakers with no reported neurological conditions signed an informed-consent sheet and received pecuniary compensation for their time. The study was approved by research ethics boards of Western University (Ontario, Canada). Due to equipment malfunction or impairments with the anaesthetic procedure three participants were excluded (1 male). Thus, 16 participants were included in this study(Naci et al., 2018).

#### Anaesthetic Procedure -LON dataset

The procedure was supervised by two anaesthesiologists and one anaesthetic nurse in the scanning room. Participants also performed an auditory target-detection task and a memory verbal recall to assess level of awareness independently from the anaesthesiologists. Additionally, an infrared camera was used to further assess level of wakefulness.

Propofol was administered intravenously using a Baxter AS50 (Singapore); stepwise increments were applied via a computer-controlled infusion pump until all three assessors agreed that Ramsay level 5 was reached (i.e. no responsiveness to visual or verbal incitements). If necessary, further manual adjustments were made to reach target concentrations of propofol which were predicted and maintained stable by a pharmacokinetic simulation software (TIVA trainer). This software also measured blood concentration levels following the Marsh 3-compartment model. The initial propofol concentration target was 0.6 μg/ml, and step-wise increments of 0.3 μg/ml were applied after which Ramsay score was assessed. This procedure was repeated until participants stopped answering to verbally and where rousable only by physical stimulation at which point data collection would begin. Oxygen titration was put in place to ensure SpO2 above 96%. The mean estimated effect site propofol concentration was 2.48 (1.82-3.14) μg/ml and propofol concentration whilst the mean plasma concentration was 2.68 (1.92-3.44). Mean total mass of propofol administered was 486.58 (1.92– 3.44). 8-minutes of RS-fMRI data was acquired.

#### Magnetic Resonance Imaging Protocol – LON dataset

A 3-tesla Siemens Trio scanner was used to acquire 256 functional volumes (Echo-planar images [EPI]). Scanning parameters were: slices=33, 25% inter-slice gap resolution 3mm isotropic; TR=2000ms; TE=30ms; flip-angle=75 degrees; matrix=64×64. Order-of-acquisition was bottom-up interleaved. The anatomical high-resolution T1 weighted images (32-channel coil 1mm isotropic voxels) were acquired using a 3D MPRAGE sequence with TA=5mins, TE =4.25ms, matrix=240×256, 9 degrees FA.

### Disorders of consciousness Dataset (DOC)

#### Patients - DOC dataset

MRI data for 23 DOC patients were collected between January 2010 and July 2015 in the Wolfson Brain Imaging Center in Addenbrookes Cambridge, UK (mean time post injury 15.75 For UWS and 16.9 for MCS). These were selected out of a bigger dataset due to their relatively intact neuroanatomy. These patients were treated and scanned at the Wolfson Brain Imaging Center, Addenbrookes hospital (Cambridge, UK). Written Informed consent was obtained from an individual that had legal responsibility on making decisions on the patient’s behalf. These participants were split into vegetative state and minimally conscious groups (n = 12 for UWS and 11 for MCS) in accordance to the diagnosis given by the attending physician at Addenbrookes hospital. Mean CRS-r score was 8.3 (standard deviation 2.03), For the UWSS group 7, (SD 1.41) and 9.75 (Sd 1.54) For the MCS group. Mean age for the UWS group was (40.16) SD 13.63; and for the MCS group (39,18, S.D 18.13). In the UWS group the aetiology was described as TBI for 3 patients, one hypoxia, one edema and the remaining participants having the pathology caused by anoxia. In the MCS group nine of the patients had a Traumatic brain injury, one a cerebral bleed and one anoxia. In the MCS group 7 were male; whilst in the UWS group 8 were male. This dataset received ethical approval from the National Research Ethics Service

#### Magnetic Resonance Imaging Protocol -DOC dataset

A varying number of functional tasks, anatomical and diffusion MRI images were taken for the DOC participants. Only the Resting-state data was used for this study. This was acquired for 10 minutes (300 volumes, TR=2s) using a siemens TRIO 3T scanner. The functional images were acquired using an echo planar sequence. Parameters include: 3×3×3.75mmm resolution, TR/TE = 2000ms/30ms, 78 degrees FA. Anatomical images T1-weighted images were acquired using a repetition time of 2300ms, TE=2.47ms, 150 slices with a cubic resolution of 1 mm.

### Preprocessing

All functional images were preprocessed in the same way using an in-house matlab script that used SPM12 functions (https://www.fil.ion.ucl.ac.uk/spm/software/spm12). After removing the first 5 scans to reach scanner equilibrium, slice-timing correction was performed (reference slice=no. 17). Volumes were realigned to the mean functional image. This process produced re-alignment parameters which were included in the time series extraction covariates. Finally, using the mean functional image, spatial normalization to an EPI-template was conducted using the function “old norm” in SPM as this yielded consistently good results. Participant-specific cerebral spinal fluid and white matter maps, used for the time series extraction (See below), were also created using an in-house Matlab (2016a) script based on SPM functions. Visual inspection of normalization to standard space was carried for all datasets. Particular attention was given to the DOC dataset because of the effect that lesions may have on spatial transformations. Due to insufficient coverage of the cerebellum in a UWS patient, these data were excluded from analyses involving the cerebellum.

#### Time Series Extraction

Denoising steps were performed in the SPM-based software CONN (17.f) (https://web.conn-toolbox.org/). Movement parameters were included as a first-level covariate. The aCompCorr algorithm regressed out CSF & White-matter signals from the time-series (using the first 5 principal components). The ART quality-assurance/motion-artifact rejection toolbox (https://www.nitrc.org/projects/artifact_detect) was also used to further clean the timeseries data. Linear de-trending and a 0.008 to 0.09 Hz band-pass filter was applied to eliminate low-frequency scanner drifts and high-frequency noise. The time-series were extracted controlling for the nuisance variables described above from the unsmoothed functional volumes to avoid artificially-induced correlations in clustered regions of interests.

#### Graph theory analysis: graph construction

Graph theory analyses were run on weighted thresholded undirected connectivity matrices (i.e., graphs). The ROIs corresponded to “nodes” and are placed on the rows and columns of a matrix; whilst the Pearson’s correlations between any two pairs of nodes were considered weighted (FC) edges and are represented by the cells in the matrix (Rubinov & Sporns, 2010). Self-connections were set to 0 and NaN values were removed to ensure graphs represented ROI-to-ROI connections.

There is no consensus regarding how to threshold connectivity matrices (Crone et al., 2014; Monti et al., 2013; Rubinov & Sporns, 2010). Usually a set of thresholds are used to ensure that results are consistent and not driven by graph topologies at specific connection densities (Hallquist & Hillary, 2018; Rubinov & Sporns, 2010; Martijn P. van den Heuvel et al., 2017). Proportional thresholding was used (e.g., top 10% of correlations). This ensures that the networks compared are of the same size, have similar properties such as node-connectivity distribution and that the density of each network was calculated relative to its size (Hallquist & Hillary, 2018; Rubinov & Sporns, 2010). There have been critiques (Hallquist & Hillary, 2018; Martijn P. van den Heuvel et al., 2017) to the use of proportional thresholding in clinical populations as baseline functional connectivity may be different compared to controls and would introduce spurious correlations in the network analysis. It is possible in this case that graph theory differences are actually driven by simple FC differences. To obviate this problem, other than controlling for dynamic FC entropy at the inferential statistic level, weighted networks were used as lower correlations would have lower values in the calculation of GTA metrics and are reported to ameliorate FC-driven GTA differences (Martijn P. van den Heuvel et al., 2017).

To further guard from the problem of the FC-driven GTA difference problems, a particularly stringent proportional threshold was used to define graphs. 5 thresholds going from 5% to 25% in 5% increases were used to test a wide-range of connection densities (Godwin et al., 2015; Monti et al., 2013). The graph theory values for each of these thresholds were then averaged to form the independent variables in inferential analyses. Only positive correlations were considered as is typical for network neuroscience due to the dubious interpretation and the preprocessing contingencies associated with negative weights (Dixon et al., 2017; Huang et al., 2020; Rubinov & Sporns, 2010).

These weighted-thresholded matrices were analysed using in-house matlab scripts which utilised functions from the brain connectivity toolbox (Rubinov & Sporns, 2010). In accordance to previous advice (Hallquist & Hillary, 2018), given how GTA results may be driven by specific parcellations (Hallquist & Hillary, 2018; Papo et al., 2016; Yao et al., 2015), the reproducibility of GTA results was tested through the use of several network definitions (see S1).

For the creation of time-varying (dynamic) connectivity matrices, a sliding-window approach was used. In accordance to previous studies (Luppi et al., 2019; Preti et al., 2017), the timeseries were split into a window composed of 24 timepoints (48 seconds) (Figure 1 in main text) which was then moved by one timepoint. The timeseries were tempered with a gaussian window to ensure that the timepoints at the edge of the windows did not have a great effect on the correlations obtained.

This procedure resulted in 122 graphs for each participant in the Cam dataset, 271 graphs for the DOC dataset and 251 for the LON dataset. The measure used upon the properties of this graph (see sample entropy section; Richman & Moorman, 2000) is meant to be relatively stable with differing number of time points.

#### Graph theory properties: definitions

Small-Worldness attempts to quantify a particular topology of self-organising complex systems(Watts & Strogatz, 1998). This particular architecture is defined by a high clustering-coefficient and a small characteristic path length.

Clustering-coefficient is defined as the fraction of neighbours of a node that are also neighbours (Humphries & Gurney, 2008; Muldoon et al., 2016; Rubinov & Sporns, 2010; Telesford et al., 2011; Watts & Strogatz, 1998) effectively operationalized as:

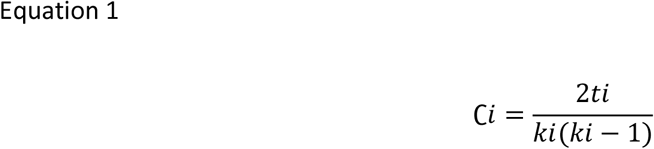

Where *t*, the number of connected triangles of node *i*, is compared to the number of connections (k) of that node. The clustering coefficient is averaged across nodes to the typical “cliquishness” of a network (Humphries & Gurney, 2008; Muldoon et al., 2016; Telesford et al., 2011; Watts & Strogatz, 1998).

Characteristic path-length is calculated as the average of the shortest distance between all pairs of nodes (Watts & Strogatz, 1998) using Dijkstra’s algorithm and is denoted as L. Small values of L indicate that information is readily available across the network(Rubinov & Sporns, 2010; Telesford et al., 2011).

It is common practice to normalise L and C to equivalent (i.e. with comparable network properties) Erdos-Renyi random networks (*Crand* & *Lrand* (Humphries & Gurney, 2008; Monti et al., 2013; Schroter et al., 2012)). This ensures that clustering coefficient and path-length rather than other network properties influence SW, and thus somewhat faithfully operationalises the original SW definition (i.e., C>>Crand & L≥Lrand (Humphries & Gurney, 2008; Watts & Strogatz, 1998)). The randomisation parameters and the number of random networks created were assessed in terms of convergence of values (i.e., recalculating with increasing values until results were consistently similar). Each Crand and Lrand were calculated from 50 random networks from a rewiring parameter of 5 (in the ranmio_und function in BCT toolbox).

The ratio between these random-network normalized values of these gives small-worldness:

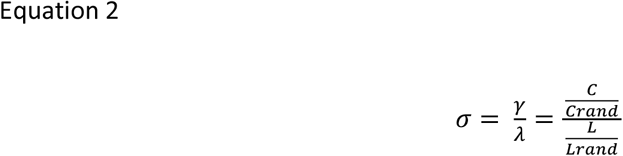

Thus, the shorter the normalised path-length (λ) and the higher the normalised clustering coefficient (γ), the higher SW-σ.

However, despite being extensively used in the literature (Lord et al., 2017; Luppi et al., 2019; Monti et al., 2013; Schroter et al., 2012), this metric has been criticized as σ is highly dependent on small variations of clustering-coefficient in the random network and is a measure that is primarily driven by clustering coefficient(Papo et al., 2016; Telesford et al., 2011). It is argued that Crand is an inappropriate normalisation model as high clustering is found in lattice networks and in fact in the original definition compares the clustering coefficient of a SW network to that of a lattice network and the path length to a random graph (Watts & Strogatz, 1998). Therefore, Telesford and colleagues (2011) (Telesford et al., 2011) suggest normalising C to the clustering coefficient of an equivalent lattice network (Clatt).

In fact, for this study the alternative function to calculate small world topology was taken from Muldoon and colleagues (Muldoon et al., 2016), which similarly to Telesford and colleague’s measure (Telesford et al., 2011), uses both lattice and random networks to normalise C and L.

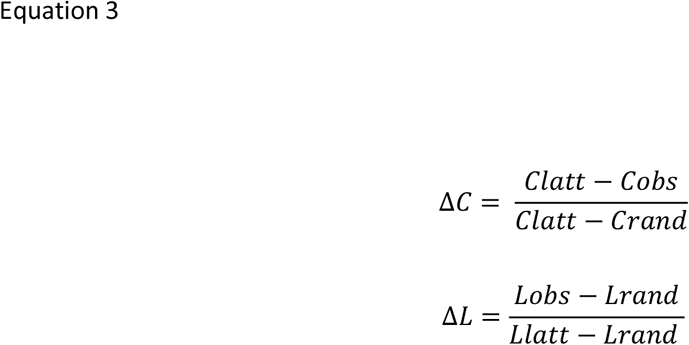

Where L and C indicate path length and Clustering Coefficient respectively of the observed network (obs), an equivalent latticed network (latt), and a random network (rand). These normalisations in turn give the SW measure which ranges from 0 to 1 (the algorithm forces values to 1 in the cases they are above this value):

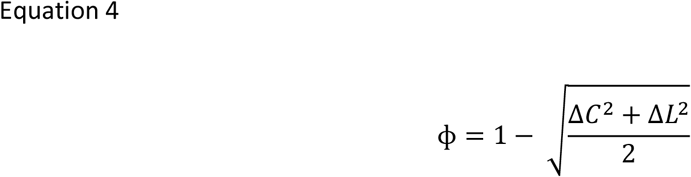

For further conceptual and statistical evaluation of the two small-world measures used in this study see supplementary material 3.

The modularity algorithm(Rubinov & Sporns, 2010) works by detecting the (computationally) optimal community structure by dividing the network into groups of nodes with maximised within group connections and minimised between group connection. Here we used the weighted version of modularity (Rubinov & Sporns, 2010).

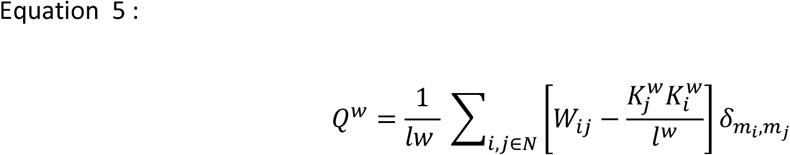

Where l is the number of links, i and j represent nodes, W the weights, and K the degree and the 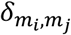 parameter is 1 if the nodes i and j are in the same module and 0 otherwise.

The participation coefficient is a measure of the richness of inter-modular connectivity of all nodes, and requires modularity to have been calculated already (Rubinov & Sporns, 2010).

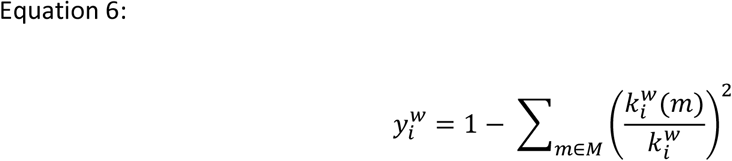

Where M is the set of modules, 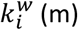 is number of weighted links between i and all nodes in module m.

All metrics were calculated across the 5 thresholded networks and the results averaged. All graph theory measures, excepting the SW propensity (Muldoon et al., 2016), were calculated using the brain connectivity toolbox (Rubinov & Sporns, 2010).

#### Sample entropy

In dynamical systems, entropy is a measure of the rate of information produced. Sample entropy was developed specifically to obviate the problem of having short and noisy timeseries which is typical of biological datasets (Delgado-Bonal & Marshak, 2019; Richman & Moorman, 2000). Sample Entropy is derived from approximate entropy, which in turn is based upon Kolomogorov complexity (Kolmogorov, 1965; Mitchell, 2011). The underlying notion being that a complex system cannot be easily described, whilst a simple system can be quickly and briefly summarized.

Sample entropy takes two timeseries segments of different lengths and compares how well each of these segments explains the rest of the timeseries (via the default Chebyshev distance measure). Sample entropy is a ratio between how well the smaller segment explains the data compared to the larger segment, and thus higher values indicating decreased self-similarity and increased complexity.

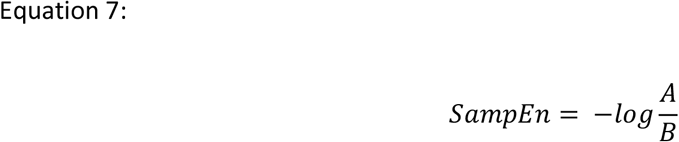

Where A is how similar the smaller timeseries segment (via Chebyshev distance) to the rest the timeseries. B is how similar the bigger timeseries segment relates to the rest of the timeseries.

The sequence lengths or timeseries lengths (max=2, min=1) were taken from a study which has looked at sample entropy of graph theory properties in functional MRI (Pedersen, Omidvarnia, Walz, Zalesky, & Jackson, 2017). Also taken from this study is the tolerance for accepting matches of similarity which was set to 0.2 times the standard deviation. The algorithm used in this paper was used in a previous study with the original creators of the Sample entropy algorithm(Richman & Moorman, 2000).

#### Inferential Analyses: Ordinal Logistic Regression

To assess the hypothesis that the dynamical complexity of graph theory properties diminished with decreasing levels of awareness Ordinal Logistic Regressions were performed using the polr function of the MASS R toolbox. This is a regression model for ordinal categorical dependent variables whilst the independent variable is continuous. This is derived from the logistic regression and ideally suited to this study for the little assumptions underlying it. Nonetheless, multicollinearity was assessed when multiple predictor variables were included and the proportional odds assumption was tested using Brants test (using package ‘brant’). The proportional odds assumption entails the model coefficients have a proportional effect on each group; I.e., “the slope” estimated between each condition (outcome variable) is the same or proportional. All tests were one-sided given the direction-specific hypotheses.

## Acknowledgements

This work was supported by grants from the UK Medical Research Council (U.1055.01.002.00001.01) [to A.M.O. and J.D.P.]; Grant from the Wellcome Trust : Clinical Research Training Fellowship to RA (grant number: 083660/Z/07/Z); The James S. McDonnell Foundation [to A.M.O. and J.D.P.]; The Canada Excellence Research Chairs program (215063) [to A.M.O.]; National Institute for Health Research (NIHR, UK); The Canadian Institute for Advanced Research (CIFAR) [to A.M.O., D.K.M. and E.A.S.]; The National Institute for Health Research (NIHR, UK), Cambridge Biomedical Research Centre and NIHR Senior Investigator Awards [to J.D.P. and D.K.M.]; The British Oxygen Professorship of the Royal College of Anaesthetists [to D.K.M.]; The Evelyn Trust, Cambridge and the EoE CLAHRC fellowship [to J.A.]; The L’Oreal-Unesco for Women in Science Excellence Research Fellowship [to L.N.]; The Stephen Erskine Fellowship, Queens’ College, University of Cambridge [to E.A.S.]; the Gates Cambridge Trust [to A.I.L.] and the Cambridge Trust [to P.C. and L.S.]. The research was also supported by the NIHR Brain Injury Healthcare Technology Co-operative based at Cambridge University Hospitals NHS Foundation Trust and University of Cambridge. Computing infrastructure at the Wolfson Brain Imaging Centre (WBIC-HPHI) was funded by the MRC research infrastructure award (MR/M009041/1). We would like to thank Victoria Lupson and the staff in the Wolfson Brain Imaging Centre (WBIC) at Addenbrooke’s Hospital for their assistance in scanning. We would also like to thank all the participants for their contribution to this study.

## Code Availability

The Coon toolbox is available to download from http://www.nitrc.org/projects/conn. The brain connectivity toolbox, which was used for the graph theory measures can be obtained gratuitously online (https://sites.google.com/site/bctnet/). Sigma was calculated fusing the BCT functions and following the procedure outlined in Humphries et al., 2008. The small world propensity measure code (PHI) can be obtained from https://complexsystemsupenn.com/codedata. The sample entropy function is freely available (https://physionet.org/content/sampen/1.0.0/).

## Author contributions

P.C. and E.A.S designed the study. P.C. carried out the analyses with advice from L.R.B.S. and A.I.L. P.F., R.A., A.M.O., L.N., G.B.W., J.A., J.D.P., D.K.M. and E.A.S. designed the original studies and collected the data. P.C. and E.A.S wrote the paper. All co-authors had the opportunity to provide commentary and feedback.

