## Supplementary Materials for "Network dynamics scale with levels of awareness"

### SUPPLEMENTARY MATERIALS – NETWORK DYNAMICS SCALE WITH AWARENESS

#### Supplementary Material 1

##### Parcellation Description

To ensure results were not dependent on specific region-of-interest (ROI) definitions, timeseries were extracted using several atlases. To obtain whole brain coverage, similarly to previous studies of interest (Monti et al., 2013; Schroter et al., 2012) the Schaefer cortical parcellations (Schaefer et al., 2018) were united with the Melbourne subcortical atlas (Tian, Margulies, Breakspear, & Zalesky, 2020) and a cerebellar atlas (Ren, Guo, & Guo, 2019). These parcellations have the advantage that they all come with different granularities permitting combinations that have a reasonable degree of similarity in terms of ROI size. Additionally, these parcellations were all defined in a data driven manner using functional connectivity (Ren et al., 2019; Schaefer et al., 2018; Tian et al., 2020).

We thus combined these Cortical, Subcortical and Cerebellar atlases to form whole brain parcellations with 126 (WB126) and 553 regions (WB553).

**Table 1.** Composition of Whole Brain Parcellations (WB126 & WB553)

| Parcellations | Cortical (Schaefer et al., 2018) | Subcortical (Tian et al., 2020) | Cerebellar (Ren et al., 2019) |
| --- | --- | --- | --- |
| WB126 | 100 | 16 | 10 |
| WB553 | 400 | 54 | 99 |

**Figure 1. Whole Brain Parcellation -126 parcels** (Ren et al., 2019; Schaefer et al., 2018; Tian et al., 2020; Xia, Wang, & He, 2013)

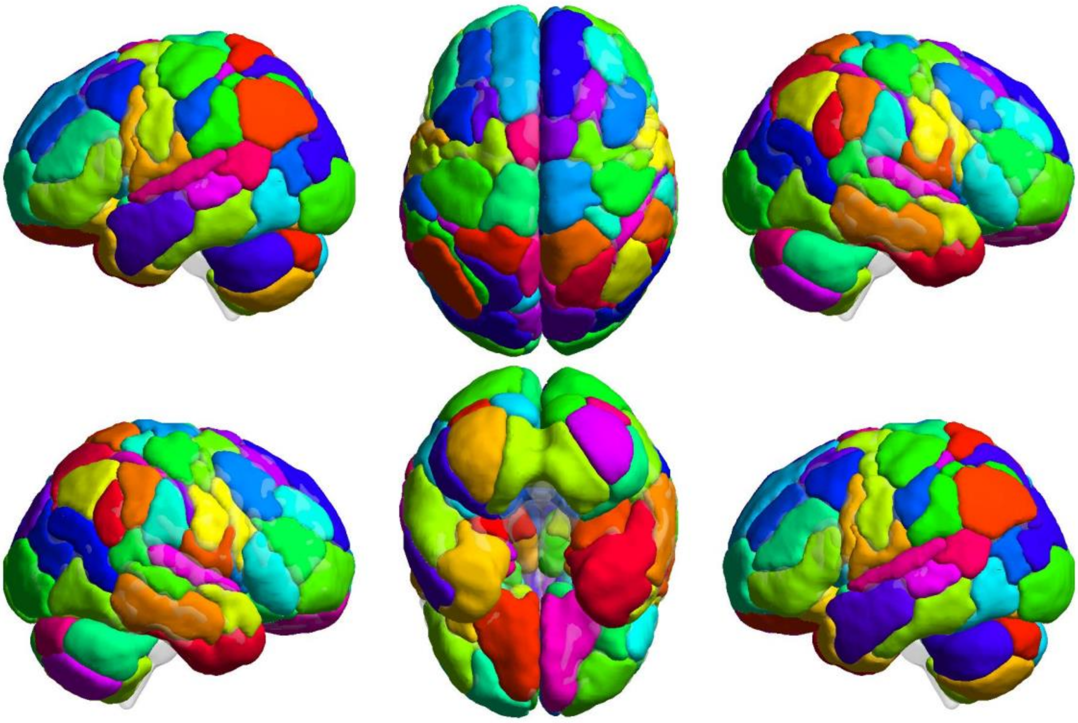

**Figure 2. Whole Brain Parcellation – 553 parcels** (Ren et al., 2019; Schaefer et al., 2018; Tian et al., 2020; Xia et al., 2013)

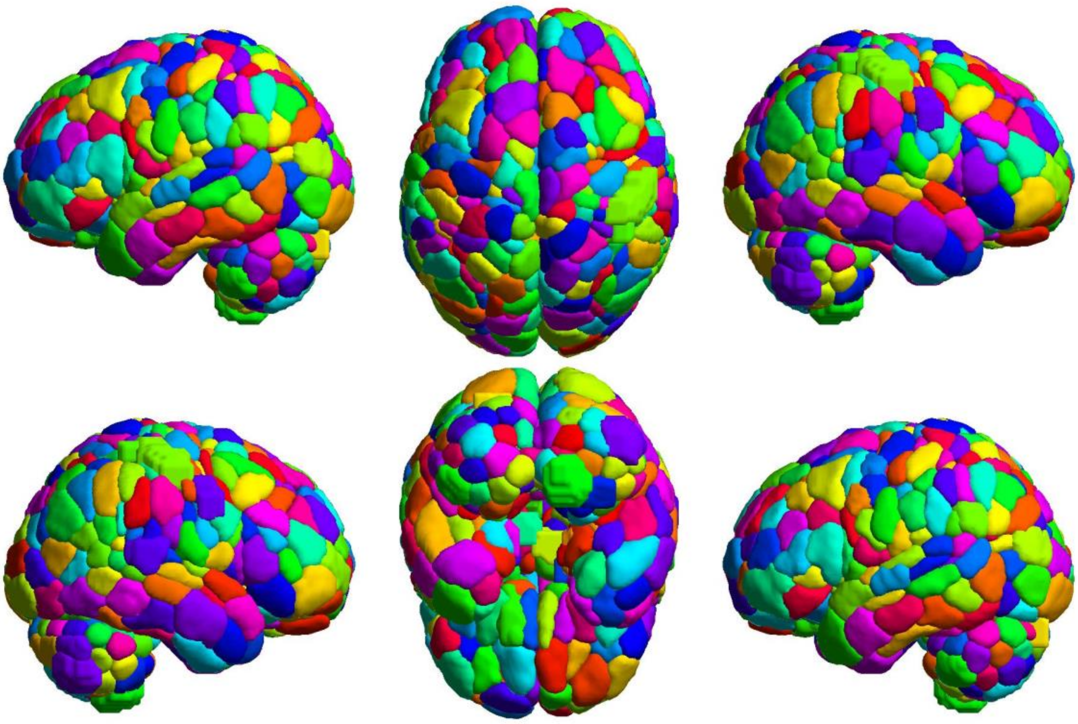

Links for download

Schaefer Cortical Atlas:

https://github.com/ThomasYeoLab/CBIG/tree/master/stable_projects/brain_parcellation/Schaefer2018_LocalGlobal

Melbourne Subcortical Atlas: https://github.com/yetianmed/subcortex/blob/master/Group-Parcellation/3T/Subcortex-Only/Tian_Subcortex_S4_3T_label.txt

Cerebellar Atlas : http://neuroguo.com/resources/

There were some overlaps in ROIs between the Cerebellar (Ren et al., 2019) and the Cortical (Schaefer et al., 2018) parcellations (154 voxels for both WB126 and WB553). These overlaps were subtracted from the cerebellar atlas in creating of the whole brain parcellations. When individual subsystems were used in any analysis, the complete original parcellations were used.

Furthermore the Automatic Anatomical Labelling Atlas (AAL) was used (Tzourio-Mazoyer et al., 2002), to further establish the stability of results and to provide results that are comparable with previous studies of interest (Luppi et al., 2019; Schroter et al., 2012). This range of parcellations were specifically used to test if results converged across different granularities and to enable exploration as to the effect of different brain structures (Telencephalon, Diencephalon and Cerebellum) on dynamic graph theory property entropy. Parcellation figures were created with BrainNet viewer (Xia et al., 2013).

**Figure 3. Automatic Anatomical Labelling Atlas (AAL) – 116 parcels.**

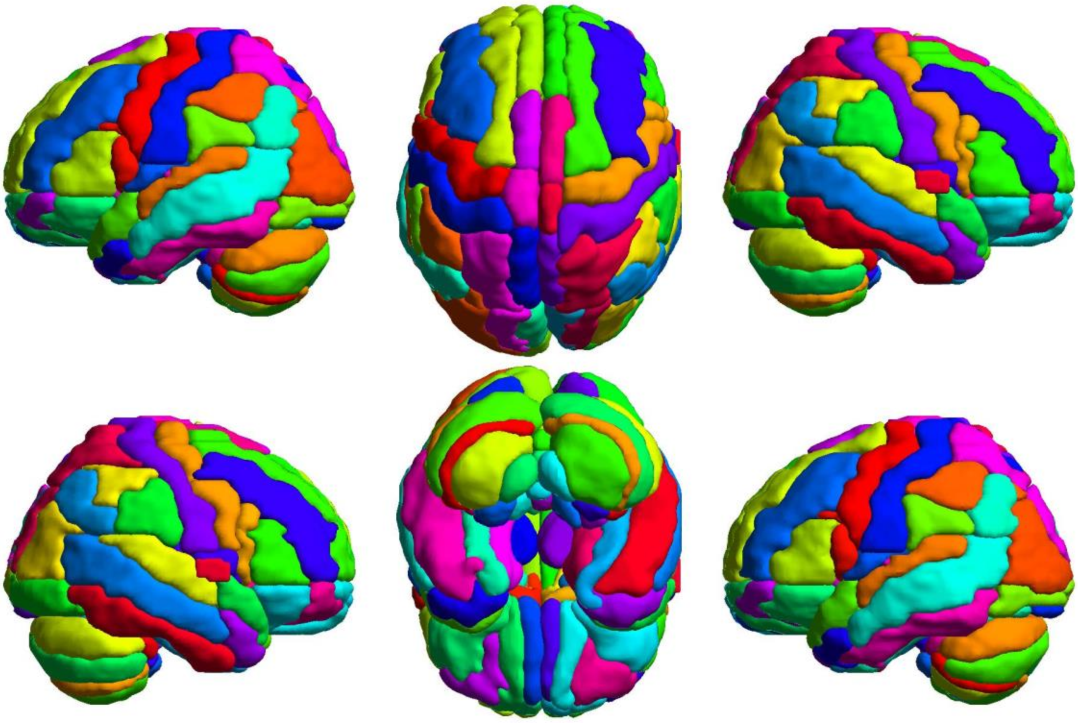

### Supplementary Material 2

**Whole brain results across different parcellations in the main and second analysis**

*Ordinal Logistic Regression statistics with Sample Entropy Of dynamic SW (PHI and Sigma) as the predictor variable.*

Predictor variables were standardized to enable comparison across analyses.

Note: dynamic sigma sample entropy was not calculated for WB553 as this was computationally intractable due to the number of connections that need to be randomised for each timepoint. This is necessary for the sigma measure to normalise values of path length and clustering coefficient (see methods section).

Brant's test (https://cran.r-project.org/web/packages/brant/brant.pdf) indicates whether the proportional odds assumption has been violated (P<0.05). Omnibus tests reported. Confidence intervals are from 2.5% to 97.5%. Outcome variables are ordered conditions (i.e. Control Awake> Sedation> MCS>UWS).

**Table 2. Main analyses (CAM-DOC datasets) – Sample entropy of SW**

| Parcellation And SW measure | SW measure | Regression Coefficients | Confidence intervals (2.5%:97.5%) | P-Value | Brant's Test |
|---|---|---|---|---|---|
| WB126 | ϕ | -1.42 | -2.04: -0.80 | 0.000003 | 0.8 |
| | **σ** | -0.91 | -1.43: -0.39 | 0.000269 | 0.45 |
| AAL | ϕ | -1.24 | -1.80: -0.68 | 0.000006 | 0.49 |
| | **σ** | -0.78 | -1.29: -0.27 | 0.001210 | 0.93 |
| WB553 | Φ | -1.12 | -1.66: -0.57 | 0.000027 | 0.75 |

**Table 3. Second Analysis (LON-DOC datasets) – Sample entropy of SW**

| Parcellation And SW measure | SW measure | Regression Coefficients | Confidence intervals (2.5%:97.5%) | P-Value | Brant's Test |
|---|---|---|---|---|---|

| | | | | | |
|---|---|---|---|---|---|
| WB126 | Φ | -0.82 | -1.37: -0.26 | 0.001932 | 0.41 |
| | **σ** | -0.81 | -1.32: -0.29 | 0.001061 | 0.4 |
| AAL | Φ | -0.85 | -1.39: -0.32 | 0.000814 | 0.19 |
| | **σ** | -0.43 | -0.93: 0.05 | 0.041476 | 0.76 |
| WB553 | Φ | -0.85 | -1.37: -0.33 | 0.000617 | 0.7 |

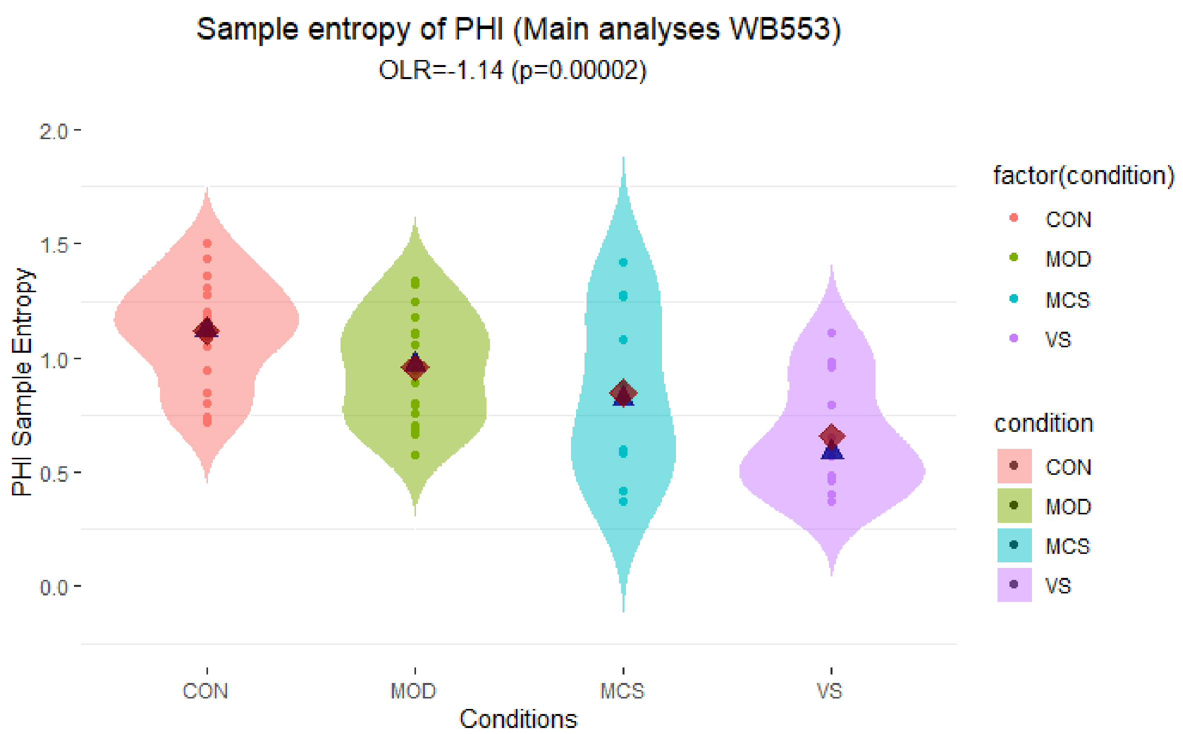


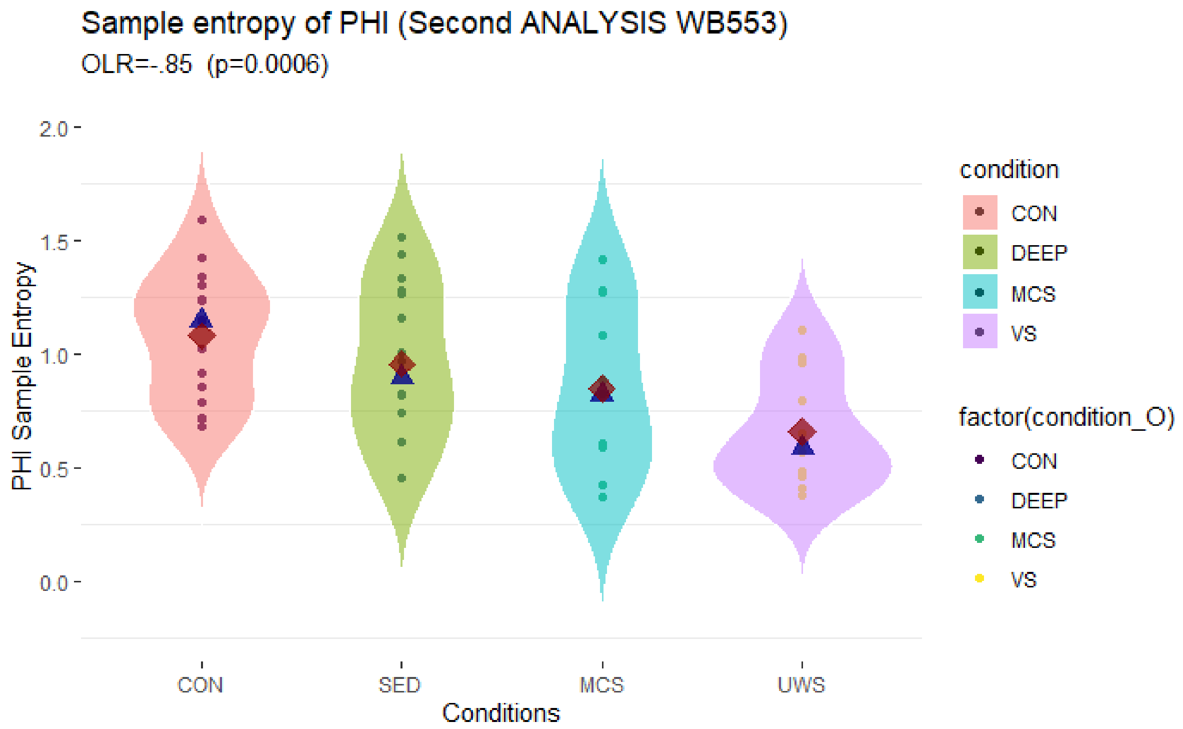


**Figure 4**. Boxplots showing ordered conditions (left to right according to presumed level of awareness: Control Awake > Sedation > Minimally Conscious State > Unresponsive Wakefulness Syndrome). PHI is a SW measure. MOD indicates moderate sedation (CAM dataset); SED indicates sedation; DEEP indicates deep anaesthesia condition (LON dataset). Y axis shows sample entropy of dynamic PHI.

These are for the WB553 parcellation. Shown are the ordinal logistic regression coefficients (OLR) with corresponding p value. Figures produced in R:ggplot2.

### Supplementary Material 3

**Matrices showing median differences of static SW measures between control awake and all other conditions.**

This supplementary section has the purpose of showing inconsistencies in static SW measures between conditions, parcellations and datasets. Static SW measures were calculated on graphs constructed using the all the timeseries available for each participant. The two static SW measures are PHI (ϕ [Muldoon, Bridgeford, & Bassett, 2016]) and Sigma ($\sigma$ [Humphries & Gurney, 2008]).

It would be expected that the two SW measures would display consistent differences between conditions and across parcellation. Additionally, if static SW was relevant to awareness, it would be expected that they would show the same direction in differences when conditions characterised by reduced awareness (i.e., Sedation, MCS, UWS) are compared to the awake control differences.

Presented are “Static SW consistency matrices”. Each cell of the matrix is the median difference between conditions. The columns represent specific subtractions (I.e. Control Awake – Sedation; Control Awake – MCS; Control Awake - UWS). The rows represent the parcellation with which the SW measures (ϕ or σ) were calculated.

Displayed are the consistency matrices for the two small worldness (Humphries & Gurney, 2008; Muldoon et al., 2016) measures calculated statically (i.e., constructing one graph using all available timepoints). Along the y-axis the parcellations and their subdivisions are indicated as well as which small world measure was used (ϕ & σ). WB126 = whole brain parcellation with 126 parcels; WB553= whole brain parcellation with 553 parcels; AAL=Automatic Labelling Atlas; CorSub = The WB553 atlas without the cerebellum (refer to supplementary material 1). Cor1= Cortex 100 parcels – Schaefer; Cor4 = Cortex 400 parcels- Schaefer; SUB = subcortex 54 parcels, Melbourne subcortical atlas. Con – Sed = median control awake – median sedation; Con- MCS = median control awake – median Minimally Conscious State; Con – UWS = median control awake - median Unresponsive Wakefulness Syndrome.

**Figure 5. Static SW consistency Matrix – Main Analysis (CAM-DOC datasets)**

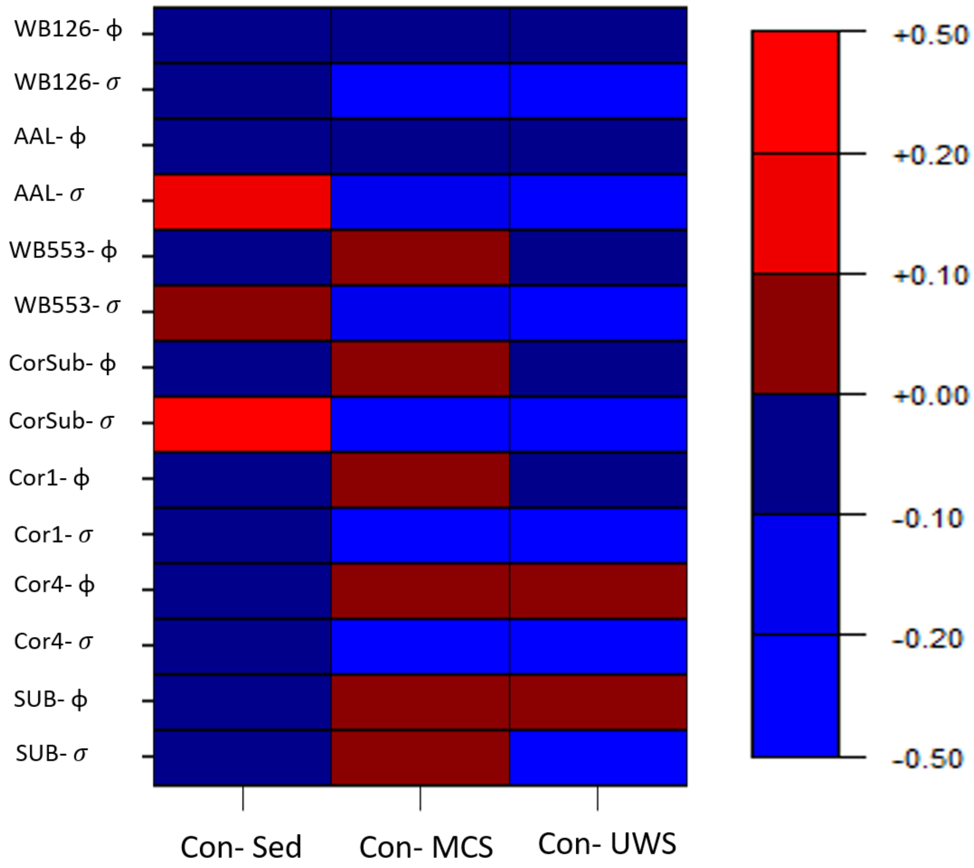

**Figure 6. Static SW consistency Matrix – Second Analysis (LON-DOC datasets)**

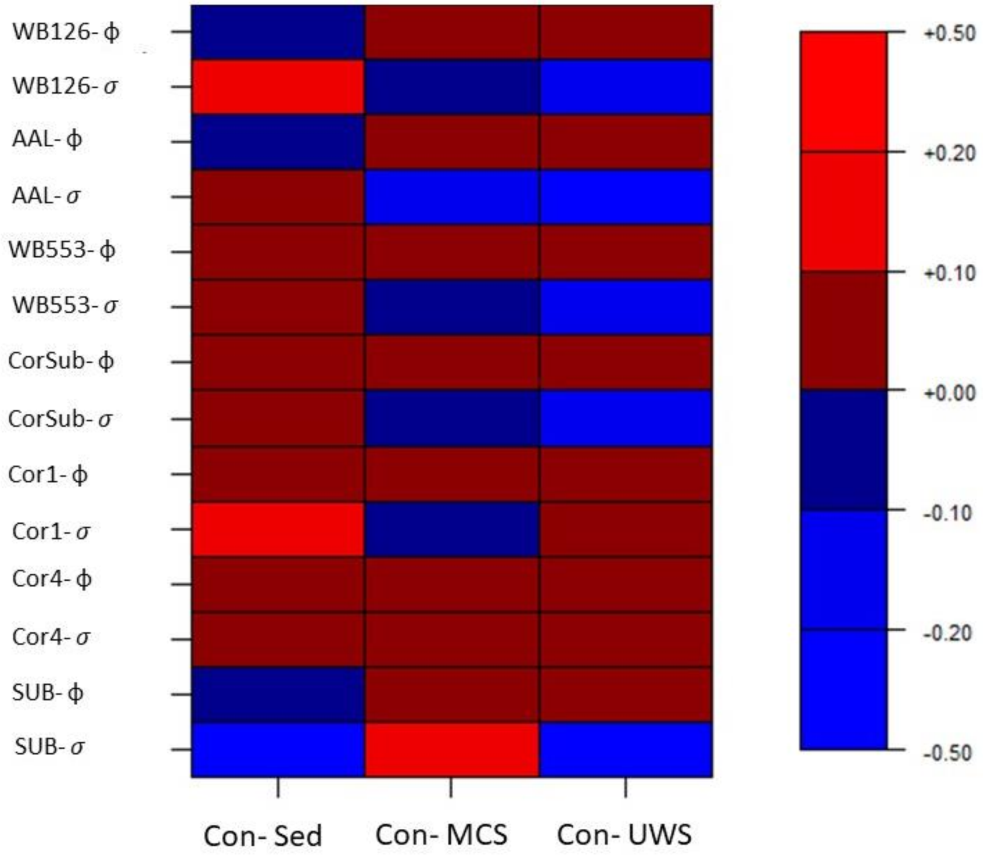


**Figure 7. Static SW measures (ϕ & $\sigma$) are not correlated with each other**

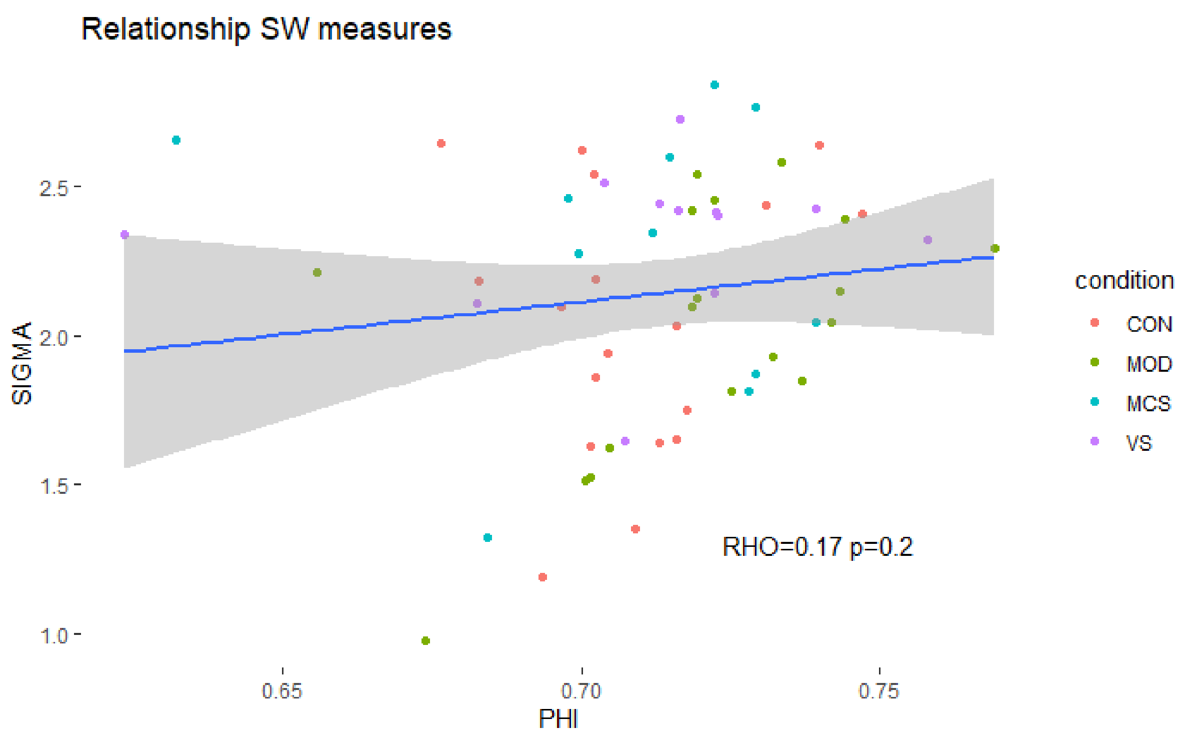


Phi was chosen as an alternative SW measure as it is not susceptible to small changes in clustering coefficient in the random networks (Papo, Zanin, Martínez, & Buldú, 2016; Telesford, Joyce, Hayasaka, Burdette, & Laurienti, 2011); ϕ Seems to be more stable over different network densities compared to sigma (Muldoon et al., 2016) and was designed to be used in weighted networks unlike the measure proposed by Telesford and colleagues (i.e., Omega [Telesford et al., 2011]). Similarly to Omega, PHI appears to be relatively stable with different numbers of nodes which is not the case with Sigma (Humphries & Gurney, 2008). Due to the use of proportional thresholding this should not constitute a confound in the present study. Furthermore, the PHI randomisation and latticisation algorithms (used to create a synthetic lattice network where neighbouring nodes are connected) reportedly maintain both degree distribution and connectivity weight distribution whilst the Brain connectivity Toolbox (BCT) which is typically used to calculate sigma (Luppi et al., 2019; Schroter et al., 2012) does so only for the degree distribution (see also methods). However, to assess convergence of results and increase comparability to previous studies; both the Sigma and the PHI measures were calculated statically and across the dynamic whole brain graphs.

#### Supplementary Material 4

**Ordinal Logistic Regressions of complexity of dynamics positive functional connectivity.**

These analyses used the same OLR method as those used for the dSW-E analyses (e.g., S2). The value of the average positive connectivity for each temporally-specific graph was calculated for each participant so as to produce a timeseries. The sample entropy was then calculated on the time series of positive average FC. (See van den Heuvel et al., [2017] or main text for rational of this analysis.)

**Table 4. Main analyses- Regression coefficients, confidence intervals, p values and Brant test p values for the OLR with dFC-E as a predictor.**

| Parcellation | Regression coefficients | Confidence intervals (2.5%:97.5%) | P-value | Brant's Test |
|---|---|---|---|---|
| WB126 | -0.49 | -0.95: -0.02 | 0.018 | 0.88 |
| AAL | -0.36 | -0.83: 0.09 | 0.058 | 0.83 |
| WB553 | -0.53 | -1.01: -0.05 | 0.013 | 0.87 |

**Table 5. Second analyses- Regression coefficients, confidence intervals, p values and Brant's test p values for the OLR with dFC-E as a predictor.**

| Parcellation | Regression coefficients | Confidence intervals (2.5%:97.5%) | P-value | Brant's test |
|---|---|---|---|---|
| WB126 | -0.68 | -1.17: -0.18 | 0.0034 | 0.26 |
| AAL | -0.60 | -1.08: -0.11 | 0.0079 | 0.45 |
| WB553 | -0.71 | -1.21: -0.21 | 0.0025 | 0.37 |

### Supplementary Material 5

**Control analyses of dSW-E with complexity of dynamic functional connectivity as a covariate.**

These analyses (similar to S2, fig 2 & 3) are ordinal logistic regressions with ordered dependent conditions as the outcome variable and both the dynamic SW sample entropy (dSW-E) and dynamic FC sample entropy (dFC-E) as continuous covariate predictors (analogous to a multiple linear regression). Therefore, each participant in each condition has both dFC-E and dSW-E values. Omnibus ("regarding all") Brant's test p-values are presented; as well as regression coefficients derived from standardised measures and predictor specific p-values.

**Table 6. MAIN analyses (CAM-DOC datasets) regression coefficients, p values and (omnibus) & Brant's test p-values**

| Parcellation | SW measure | **dSW-E** | | **dFC-E** | | Brant’s test p-value |
|---|---|---|---|---|---|---|
| | | Regression coefficient | P-value | Regression coefficient | P-value | |
| WB126 | **Φ** | -1.51 | 0.00001 | 0.14 | 0.30 | 0.86 |
| | **σ** | -0.93 | 0.0002 | -0.48 | 0.019 | 0.68 |
| AAL | **Φ** | -1.35 | 0.00001 | 0.21 | 0.22 | 0.3 |
| | **σ** | -0.77 | 0.00150 | -0.35 | 0.070 | 0.95 |
| WB553 | **Φ** | -1.17 | 0.0002 | 0.09 | 0.37 | 0.93 |

**Table 7. Second analyses (LON-DOC datasets) regression coefficients, p values and (omnibus) & Brant’s test p-values**

| Parcellation | SW measure | **dSW-E** | | **dFC-E** | | Brant’s test p-value |
|---|---|---|---|---|---|---|
| | | Regression coefficient | P-value | Regression coefficient | P-value | |
| WB126 | **Φ** | -0.59 | 0.04 | 0.3526 | 0.12 | 0.048 |
| | **σ** | -0.64 | 0.01 | -0.44 | 0.049 | 0.3 |

| | | | | | | |
|---|---|---|---|---|---|---|
| AAL | **Φ** | -0.77 | 0.01 | -0.11 | 0.36 | 0.01 |
| | **σ** | -0.28 | 0.13 | -0.52 | 0.021 | 0.59 |
| WB553 | **Φ** | -0.65 | 0.02 | -0.31 | 0.16 | 0.24 |

Note: Brant's test is significant in the second analysis for WB126 PHI and AAL PHI. This suggests that the proportional odds assumption is violated, meaning that the model coefficients are not proportional among outcome variables.

### Supplementary Material 6

**Subsystem (I.e., cortex, subcortex, cerebellum) ordinal logistic regression results with dSW-E as a predictor.**

Presented are tables of ordinal logistic regressions with the dSW-E (ϕ) of the cortex (two granularities: 100 & 400 Parcels), subcortex and the cerebellum (refer to supplementary material 1). The cerebellar results are only presented for the main analyses as the second analysis did not have sufficient brain coverage during data collection.

PHI is presented exclusively as this is the most stable measure and produced the highest effect sizes in previous analyses, and is the most computationally viable measure. Subcortical and cerebellar parcellations could not have an alternative granularity parcellation as too few nodes may cause problems in graph theory analyses (Rubinov & Sporns, 2010). dSW-E values were standardised before being inserted into the ordinal logistic regression. Presented are the resulting regression coefficients, confidence intervals (2.5%:97.5%), p values and omnibus Brant's test p-values.

**Table 8. Main subsystem analysis (CAM-DOC) regression coefficients, confidence intervals, p-values, Brant's test. dSW-E (ϕ)**

| Parcellation | Regression coefficient | Confidence intervals (2.5%:97.5%) | P-value | Brant's test |
|---|---|---|---|---|
| Cortex-100 | -1.30 | -1.89: -0.71 | 0.000006 | 0.16 |
| Cortex-400 | -1.00 | -1.61: -0.48 | 0.00017 | 0.61 |
| Subcortex | -1.94 | -2.79: -1.10 | 0.000003 | 0.76 |
| Cerebellum | -0.52 | -1.06: 0.016 | 0.02865 | 0.21 |

**Table 9. Second subsystem analysis (CAM-DOC) regression coefficients, confidence intervals, p-value, Brant's test. dSW-E (ϕ)**

| Parcellation | Regression coefficient | Confidence intervals (2.5%:97.5%) | P-value | Brant's test |
|---|---|---|---|---|
| Cortex-100 | -0.87 | -1.42: -0.32 | 0.00087 | 0.26 |

| | | | | |
|---|---|---|---|---|
| Cortex-400 | -0.35 | -0.84: 0.14 | 0.08158 | 0.53 |
| Subcortex | -2.08 | -3.08: -1.08 | 0.00002 | 0.53 |
| Cerebellum | NA | NA | NA | NA |

Note: there was insufficient coverage of cerebellum in the LON dataset.

### Supplementary Material 7

**Control analyses of subsystem dSW-E complexity with dynamic functional connectivity entropy as a covariate and subsystem odds ratio comparison.**

Presented are tables with ordinal logistic regression results for the cortex (100 & 400 parcels), the subcortex and the cerebellum (the latter only for the main analysis as the LON dataset did not have sufficient brain coverage). The object of this analyses is to investigate whether the dSW-E remains significant despite controlling for the dynamic FC sample entropy from which is derived, as this may explain the relevant variance and change due interpretations (van den Heuvel et al., 2017). All values were standardised before being inserted into the ordinal logistic regressions. Presented are the resulting regression coefficients and p-values for each predictor (dSW-E & dFC-E) and omnibus Brant's test p-values.

**Table 10. Main analyses- subsystem ordinal logistic regression results with dynamic functional connectivity entropy as a covariate to dSW-E.**

| Parcellation | SW measure | **dSW-E** | **dFC-E** | Brant's test p-value |
|---|---|---|---|---|

| | | Regression coefficient | P-value | Regression coefficient | P-value | |
|---|---|---|---|---|---|---|
| Cortex-100 | ϕ | -1.25 | 0.0001 | -0.10 | 0.36 | 0.04 |
| Cortex-400 | ϕ | -0.99 | 0.0008 | -0.02 | 0.45 | 0.72 |
| Subcortex | ϕ | -1.83 | 0.00001 | -0.47 | 0.03 | 0.51 |
| Cerebellum | ϕ | -0.46 | 0.055 | -0.13 | 0.29 | 0.34 |

**Table 11. Second Analyses- subsystem ordinal logistic regression results with dynamic functional connectivity entropy as a covariate to dSW-E.**

| Parcellation | SW measure | **dSW-E** | | **dFC-E** | | Brant's test p-value |
|---|---|---|---|---|---|---|
| | | Regression coefficient | P-value | Regression coefficient | P-value | |
| Cortex-100 | ϕ | -0.74 | 0.0085 | -0.27 | 0.1744 | 0.01 |
| Cortex-400 | ϕ | -0.07 | 0.401 | -0.43 | 0.0083 | 0.53 |
| Subcortex | ϕ | -1.94 | 0.00009 | -0.45 | 0.0506 | 0.21 |

**Note**: cerebellum was not analysed for the second analysis due to incomplete MRI coverage

### SUPPLEMENTARY MATERIAL 8

**Analyses of the complexity of dynamic small worldness when subsystems are inserted as covariates in the same ordinal logistic regression**

To investigate the difference in predictive power the dSW-E of different subsystems, we inserted these values for all available subsystems into the same ordinal logistic regression as covariates. We standardized all values, and obtained odds ratios, which may be considered a measure of effect size. As shown below in the second analyses (mirroring the primary analysis presented in the main text), the subcortex displays a strong predictive power compared to the cortex (100 parcels). To aid interpretability the dependent variable was ordered in an inverse manner compared to previous analyses (i.e., UWS>MCS>Sedation>Control Awake).

**Table 12. Main Analyses- comparison of standardized dSW-E odds ratios of subsystems when inserted into the same model**

| | Measure | Odds ratio | C.I. (2.5:97.5) | P value | Brant's test p-value |
|---|---|---|---|---|---|
| Cortex | dSW-E (PHI) | 2.81 | 1.50: 5.52 | 0.000838 | 0.36 |
| Subcortex | dSW-E (PHI) | 6.34 | 2.60: 17.71 | 0.000071 | |
| Cerebellum | dSW-E (PHI) | 1.26 | 0.70: 2.32 | 0.021628 | |

**Table 13. Second Analyses- Comparison of standardized dSW-E odds ratios of subsystems when inserted into the same model**

| | Measure | Odds ratio | C.I. (2.5:97.5) | P value | Brant's test p-value |
|---|---|---|---|---|---|
| Cortex | dSW-E (PHI) | 1.64 | 0.90:3.07 | 0.052 | 0.64 |
| Subcortex | dSW-E (PHI) | 7.53 | 2.93:22.50 | 0.00004 | |

### Supplementary Material 9

**Analyses of the complexity of dynamic modularity and participation coefficient when subsystems are inserted as covariates in the same ordinal logistic regression**

**Table 14. Main Analyses- Comparison of standardized dQ-E (modularity) odds ratio of subsystems when inserted into the same model**

| | Measure | Odds ratio | C.I. (2.5:97.5) | P value | Brant's test p-value |
|---|---|---|---|---|---|
| Cortex | dQ-E | 1.52 | 0.86: 2.75 | 0.073 | 0.09 |
| Subcortex | dQ-E | 2.15 | 1.10: 4.42 | 0.139 | |
| Cerebellum | dQ-E | 1.61 | 0.85: 3.07 | 0.069 | |

**Table 15. Main Analyses- Comparison of standardized dPC-E (participation coefficient) odds ratios of subsystems when inserted into the same model**

| | Measure | Odds ratio | C.I. (2.5:97.5) | p value | Brant's test p-value |
|---|---|---|---|---|---|
| Cortex | dPC-E | 1.09 | 0.63: 1.90 | 0.36689 | 0.5 |
| Subcortex | dPC-E | 2.14 | 1.08: 4.44 | 0.01639 | |
| Cerebellum | dPC-E | 2.77 | 1.46: 5.71 | 0.00156 | |

**Table 16. Second Analyses- Comparison of standardized dQ-E (modularity) odds ratio of subsystems when inserted into the same model**

| | Measure | Odds ratio | C.I. (2.5:97.5) | P value | Brant’s test p-value |
|---|---|---|---|---|---|
| Cortex | dQ-E | 1.14 | 0.65: 2.04 | 0.3204 | 0.55 |
| Subcortex | dQ-E | 2.58 | 1.43: 4.89 | 0.0011 | |

**Table 17. Second Analyses- Comparison of standardized dPC-E (participation coefficient) odds ratios of subsystems when inserted into the same model**

| | Measure | Odds ratio | C.I. (2.5:97.5) | P value | Brant’s test P-value |
|---|---|---|---|---|---|
| Cortex | dPC-E | 0.98 | 1.24: 4.78 | 0.4812 | 0.02 |
| Subcortex | dPC-E | 2.32 | 0.55: 1.72 | 0.0063 | |
